# TREM1^+^ regulatory myeloid cells expand in steatohepatitis-HCC and associate with poor prognosis and therapeutic resistance to immune checkpoint blockade

**DOI:** 10.1101/2022.11.09.515839

**Authors:** Julie Giraud, Domitille Chalopin, Eloïse Ramel, Thomas Boyer, Atika Zouine, Marie-Alix Derieppe, Nicolas Larmonier, Olivier Adotevi, Brigitte Le Bail, Jean-Frédéric Blanc, Laurence Chiche, Macha Nikolski, Maya Saleh

## Abstract

Hepatocellular carcinoma (HCC) is an inflammation-associated cancer arising from viral and non-viral etiologies. Immune checkpoint blockade primarily benefits patients with viral HCC. Expansion of suppressive myeloid cells is a hallmark of chronic inflammation and cancer, but their heterogeneity in HCC is not fully resolved and might underlie immunotherapy resistance in the steatohepatitis setting. Here, we present a high resolution atlas of hepatic innate immune cells from patients with HCC that unravels a steatohepatitis contexture characterized by the emergence of high entropy myeloid cell states and myeloid-biased NK cell differentiation. We identify a discrete population of tumor-infiltrating myeloid cells, predominant in the steatohepatitis setting, that expresses a variety of myeloid lineage-affiliated genes, including granulocyte, macrophage and dendritic cell features, and can be identified in HCC tumors based on selective dual expression of *TREM1* and *CD163*. Functional characterization reveals that TREM1^+^ CD163^+^ myeloid cells highly express TGFβ and IL-13RA, localize to HCC fibrotic lesions, and potently suppress T cell effector functions *ex vivo*, a function further potentiated by TREM1 engagement. We refer to this population as TREM1^+^ CD163^+^ regulatory myeloid cells (TREM1^+^CD163^+^ M_reg_). Deconvolution analyses in large cohorts of patients with HCC and other solid tumors reveals that the density of TREM1^+^ CD163^+^ M_reg_ increases in advanced stages, associates with poor prognosis, and therapeutic resistance to PD-1 blockade. Our data support myeloid subset-targeted immunotherapies to treat HCC and identify TREM1 as a therapeutic target.

**HIGHLIGHTS:**

- Atlas of hepatic innate immune cells (100,000 transcriptomes) from patients with HCC
- Core signatures to identify, discriminate and localize innate lymphoid and myeloid cells
- A population of TREM1^+^CD163^+^ myeloid cells, referred to as TREM1^+^CD163^+^ M_reg_, expands in steatohepatitis HCC
- TREM1^+^CD163^+^ M_reg_ express granulocyte- and macrophage/dendritic cell-lineage genes
- TREM1^+^CD163^+^ M_reg_ potently suppress T cell effector functions, which is potentiated by TREM1 engagement by cognate ligands
- TREM1^+^CD163^+^ M_reg_ produce high levels of TGFβ and populate fibrotic lesions
- The density of TREM1^+^CD163^+^ M_reg_ increases in advanced HCC and associate with poor patient survival
- The density of TREM1^+^CD163^+^ M_reg_ associates with resistance to immune checkpoint blockade in other solid tumors

## INTRODUCTION

Hepatocellular carcinoma (HCC) ranks among the most common malignancies worldwide, with a rising incidence in the Western World. Despite well-known risk factors, i.e. chronic viral infection with hepatitis B virus (HBV) primarily in Asia and HCV in western countries, excessive alcohol consumption and the metabolic syndrome-associated non-alcoholic steatohepatitis (NASH), HCC is diagnosed late in most patients (Llovet et al., 2021). While several systemic treatment options have been approved for advanced HCC, e.g., sorafenib and other tyrosine kinase inhibitors, they all provide a small clinical benefit (Finn et al., 2020). The landscape of clinical trials for HCC treatment has recently shifted to the field of immunotherapy. The portfolio of therapeutic options now includes the immune checkpoint inhibitors (ICI) nivolumab and pembrolizumab, and in 2020, the FDA approved the first combination therapies (atezolizumab/bevacizumab and nivolumab/ipilimumab) in HCC (Finn *et al*., 2020). However, despite significant therapeutic advance with ICI, ∼75% of patients do not respond to these immunotherapies for unclear reasons (Giraud et al., 2021). More recently, a meta-analysis of three randomized phase III clinical trials administering ICI to patients with advanced HCC showed a superior efficacy of immunotherapies in virally-infected patients compared to NASH-affected patients with HCC (Pfister et al., 2021). This suggests that the tumor microenvironment (TME) of HCC is an important determinant of therapeutic success and highlight the urgent need to further explore human liver-specific immunity towards the identification of theranostic immune biomarkers for patients stratification and novel immunotherapies.

Myeloid cells are a main immune infiltrate in several solid tumors and considered as an impediment to all cancer therapeutic modalities, particularly immunotherapy. Thus, modulating their recruitment, differentiation or functions is being actively pursued as a therapeutic option (Goswami et al., 2022). However, their indiscriminate depletion has failed to improve cancer patient overall survival, indicating that a better characterization of the deleterious myeloid subsets is required for a targeted approach. Myeloid cells encompass mononuclear phagocytes (MNP) (including monocytes, macrophages and dendritic cells [DC]) and granulocytes, which exhibit remarkable heterogeneity and divergent functions according to ontogeny, tissue microenvironment, proximal inflammatory and metabolic signals and trained immunity elicited epigenetic reprogramming. The advent of single cell and tracing technologies have unraveled the complexity of myeloid states, warranting context-specific characterization of deleterious immunosuppressive and pro-tumoral subsets to direct targeted therapies.

While the cellular landscape of HCC has been previously described using single cell RNA sequencing (scRNA-seq) and mass cytometry (Ma et al., 2019; Ma et al., 2021; Sharma et al., 2020; Song et al., 2020; Sun et al., 2021; Zhang et al., 2019; Zheng et al., 2017), the impact of etiology on the diversity of the innate immune compartment, particularly myeloid cell states, has not been fully characterized. Here, we implemented scRNA-seq on purified hepatic innate immune cells freshly isolated from tumoral and juxta-tumoral tissues from patients with HCC of different etiologies, and performed spatial transcriptomics (stRNA-seq) to map their localization. We unravel a steatohepatitis-associated contexture characterized by enhanced myeloid biased NK cell differentiation and the emergence of entropic myeloid cell states. Among the myeloid populations, we identify three *THBS1*+ myeloid cell subsets with features previously linked to a suppressive state (Condamine et al., 2016). Among these, one discrete population was abundant in the tumor of patients with steatohepatitic HCC but absent in patients with viral HCC in our cohort. We show that this population expresses a variety of myeloid lineage-affiliated genes, including granulocyte, macrophage and dendritic cell features, and can be identified in HCC tumors based on selective dual expression of TREM1 and CD163. Through functional characterization, we demonstrate that TREM1^+^CD163^+^ myeloid cells are potent suppressors of T cell effector functions *ex vivo*, a function further potentiated by TREM1 engagement. Our results reveal that TREM1^+^CD163^+^ M_reg_ express high levels of TGFβ and IL-13RA and are spatially enriched at HCC fibrotic lesions in close association with FAP^+^ cancer-associated fibroblasts. Deconvolution analyses in large cohorts of patients with HCC reveals that the intra-tumoral density of TREM1^+^CD163^+^ M_reg_ population increases in advanced stages and associates with poor prognosis according to different molecular classifications. In a separate cohort of patients with renal cell carcinoma, the density of TREM1^+^CD163^+^ M_reg_, but not that of any other MNP subset characterized in our study, also associates with therapeutic resistance to PD-1 blockade. Our results elucidate this population as a biomarker of severe HCC and a therapeutic target in patients resistant to immunotherapy.

## RESULTS

### The hepatic innate immune landscape in HCC

To characterize the innate immune cellular landscape of the liver from patients with HCC, we implemented scRNA-seq (10x Genomics) on FACS-sorted CD45^+^ panTCRαβ^-^ CD19^-^ innate immune cells freshly isolated from tumors (HCC) and adjacent non-tumoral (NT) liver sections. In parallel, we applied spatial transcriptomics (ST; Visium^®^, 10x Genomics) on frozen tissue sections from the same patients (Figure 1a). We included 10 HCC patients with different etiologies (HCV (n=3), obesity (n=5), excessive alcohol consumption (n=3)) (Figure S1a and Table S1). NASH (n=3) and cirrhotic lesions (n=3) were confirmed in HCC and NT tissue sections by hematoxylin eosin saffron and trichrome staining (Figure S1b). For the scRNA-seq, for each of the 20 fresh samples, 15,000 innate immune viable cells were loaded on the 10x chip. Following putative doublet removal and exclusion of stressed or dead cells (Figure S1c), we analyzed the transcriptomes of ∼96,000 single innate immune cells (∼2,300 genes/cell) across 10 HCC (∼46,000 cells) and 10 NT (∼50,000 cells) samples. Following data processing using the SEURAT pipeline, 22 Louvain clusters were identified (Figure 1b), containing cells derived from all patients and tissue sites (Figure 1c and Table S2). Using a scoring method based on published signatures retrieved from Panglao DB (Table S3), combined with the expression of discriminatory features (Figure 1d and Table S4) and canonical markers (Figure 1e), we identified the major hepatic innate immune cell populations (Figure 1f). In total, we identified 11 innate lymphoid clusters (∼68,000 cells), encompassing one innate lymphoid cell (ILC) cluster (c20) and 10 clusters of NK cells. These included one cycling NK cluster (c16), two NK1 clusters (c0, 2) representing *CD56*^low^ *CD16*^+^ cytotoxic NK cells, four NK2 clusters (c1, 4, 6, 9) representing inflammatory *CD56*^high^ *CD16*^-^ NK cells, two mixed NK3/γδ T cell clusters (c3, 15), and one small NK4 cluster (c13). In addition, we identified 10 myeloid clusters (∼28,000 cells) including four monocyte/macrophage (Mo/Mac) clusters (c5, 7, 10, 14), two conventional DC clusters (c11, 17), one plasmacytoid DC (pDC) cluster (c19) and three granulocyte clusters (neutrophils (c12), mast cells (c18) and basophils (c21)). Besides the expected myeloid cells, our analysis revealed a population of cells found primarily on the myeloid side of the Louvain graphical representation (c8, green) (Figure 1f), expressing both lymphoid (*IL7R, GNLY, KLRF1, KLRD1*, *GZMH/K/B, IFNG*) and myeloid genes (*LYZ*, *HLA- DRA/B1, S100A8/9, SAT1, CST3*) (Figure 1d and Figure S2). We refer to this population as myeloid NK (MyeNK), as characterized below. Next, to assess the tumoral distribution of the identified innate immune cell subsets, we quantified cluster frequencies in HCC compared to NT. This analysis revealed significant intra-tumoral depletion of NK2 cells and enrichment of Mo/Mac subsets (Figure 1g, h), and such TME remodeling is exacerbated with cancer progression (Figure 1i). Altogether, our results define the landscape of innate immune cells in HCC, confirm significant myeloid cell infiltration in tumors and unravel a distinct population of cells with the unique property of dually expressing innate lymphoid and myeloid genes.

**Figure 1.**
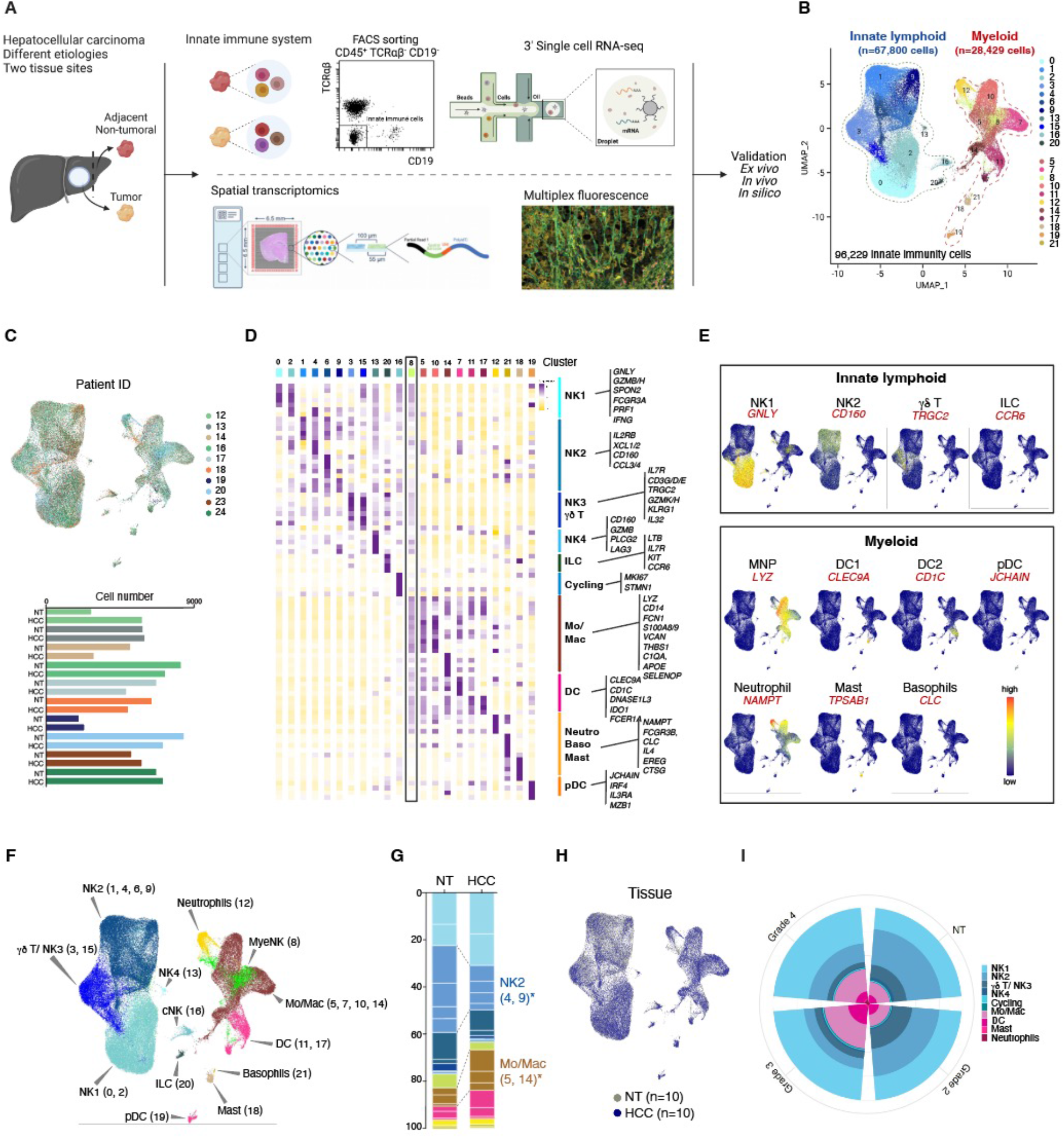
The innate immunity landscape of the liver of patients with hepatocellular carcinoma (HCC) (A) Multi-omics characterization of the hepatic innate immune cell landscape of human patients with hepatocellular carcinoma (HCC) using scRNA-seq and stRNA-seq of tumors and adjacent NT of 10 HCC patients with different etiologies. (B) Louvain clustering of 96,229 scRNA-seq transcriptomes of hepatic innate immune cells in two tissue sites from 10 HCC patients identifies 22 main clusters in the liver. (C) UMAP visualization colored according to patient ID and their respective cell number recovery according to tissue. (D) Heatmap representing average expression of discriminating genes for each cluster. (E) Feature Plots representing the expression of innate immune cells canonical markers. (F) Louvain clusters shown in (B) colored based on cell type annotation. (G) Main cluster frequencies in HCC and NT. *p<0.05 multiple Wilcoxon tests. (H) UMAP colored by tissue site, HCC (blue) and NT (grey). (I) Circular plot representing cluster frequencies in NT and HCC of different tumor grades.

### Phenotypic and functional characterization of myeloid NK cells

NK cells expressing myeloid genes have been previously described in mice as precursors to mature NK (pre-mNK) (reviewed in (Guimont-Desrochers and Lesage, 2013)). In addition, both mature NK (Theurich et al., 2017) and γδ T (Howard et al., 2017) cells can acquire myeloid properties in pathological conditions, notably in obesity (Theurich *et al*., 2017). To unravel the identity of cells in c8, we first selected genes that clearly distinguish lymphoid from myeloid cell subsets in our data, namely *NKG7, KLRD1, LYZ* and *HLA-DRA* (Figure S2a). We next ascertained dual expression of these NK/lymphoid and myeloid features on a per cell basis. This analysis showed that the majority of cells co-expressing these genes belonged to c8 (Figure S2b). Cell trajectory analysis using Slingshot revealed that c8 occupied a central position along the Pseudotime between lymphoid and myeloid lineages (Figure S2c). To distinguish among the innate lymphoid subpopulations, we next tested two signatures capable of distinguishing NK cells from ILCs (Heinrich et al., 2021) or γδ T cells (*CD3D*, *CD3E*, *TRDC* and *TRGC2*) (Pizzolato et al., 2019) (Table S5). Cells in c8 scored highest for the NK gene signature (Figure S2d). Taken their segregation with the myeloid transcriptomes (Figure 1f), we posited that c8 is a population of myeloid-biased NK cells. Phenotypically, they were CD56^low^ (*NCAM1*) CD16^+^ (*FCGR3A*), expressed cytotoxic factors (*GZMB, GZMK, IFNG*), NK activating receptors (*CD160*) but low levels of markers associated with tissue homing (*EOMES, ITGAE*) (Figure S2e). This phenotype suggests that MyeNK cells are mature cytotoxic effectors. To confirm this, we conducted computational tests and functional validation. Computationally, we estimated differentiation potency by calculating entropy rates (or signaling promiscuity), which is useful to distinguish precursor states (high entropy rates) from fully differentiated cells (low entropy rates). This analysis revealed that, unlike cycling NK (cNK) cells that displayed high entropy rates, MyeNK cells had the lowest entropy rates compared to all other innate immune cells analyzed, indicative of a differentiated state (Figure S2f). For functional validation, we first confirmed the detection of these cells in HCC surgical resections by multiplex immunofluorescence (IF) e.g. cells co-expressing GNLY and HLA-DR, and flow cytometry (FACS) e.g. cells co-expressing KLRD1 and HLA-DR (Figure S2g, h). Notably, compared to MNPs, HLA-DR was expressed at intermediate levels on MyeNK cells both at the transcript (Figure S2b, bottom) and protein (Figure S2h) levels. We next FACS-isolated them based on the co-expression of KLRD1 and HLA-DR (gated on CD45^+^ CD19^-^ panTCRαβ^-^, panTCRγδ^-^ cells) and assessed their lytic capacity in an *ex vivo* killing assay. These experiments showed that similar to NK cells, MyeNK cells responded to IL-2+IL-15 and were highly cytotoxic (Figure S2i, j). In addition, as for other NK cells, MyeNK cells were found at lower proportions in HCC compared to NT, representing 3,3% and 5,9% of all hepatic innate immune cells, respectively (Figure 1g). Re-clustering of the MNP transcriptomes unraveled two clusters of MyeNK cells (c2, 5) (Figure 2a and S2k, l). c2 originated mostly from one patient, whereas c5 contained cells from all patients (Table S2). MyeNK cells (c5) were poorly represented in the tumor (Figure S2k), but increased in the juxta-tumoral tissue of obese- and/or NASH-affected patients with HCC (Figure S2l). Altogether, our results identify a myeloid-biased NK cell population that is expanded in the liver of obese/NASH-affected patients.

**Figure 2.**
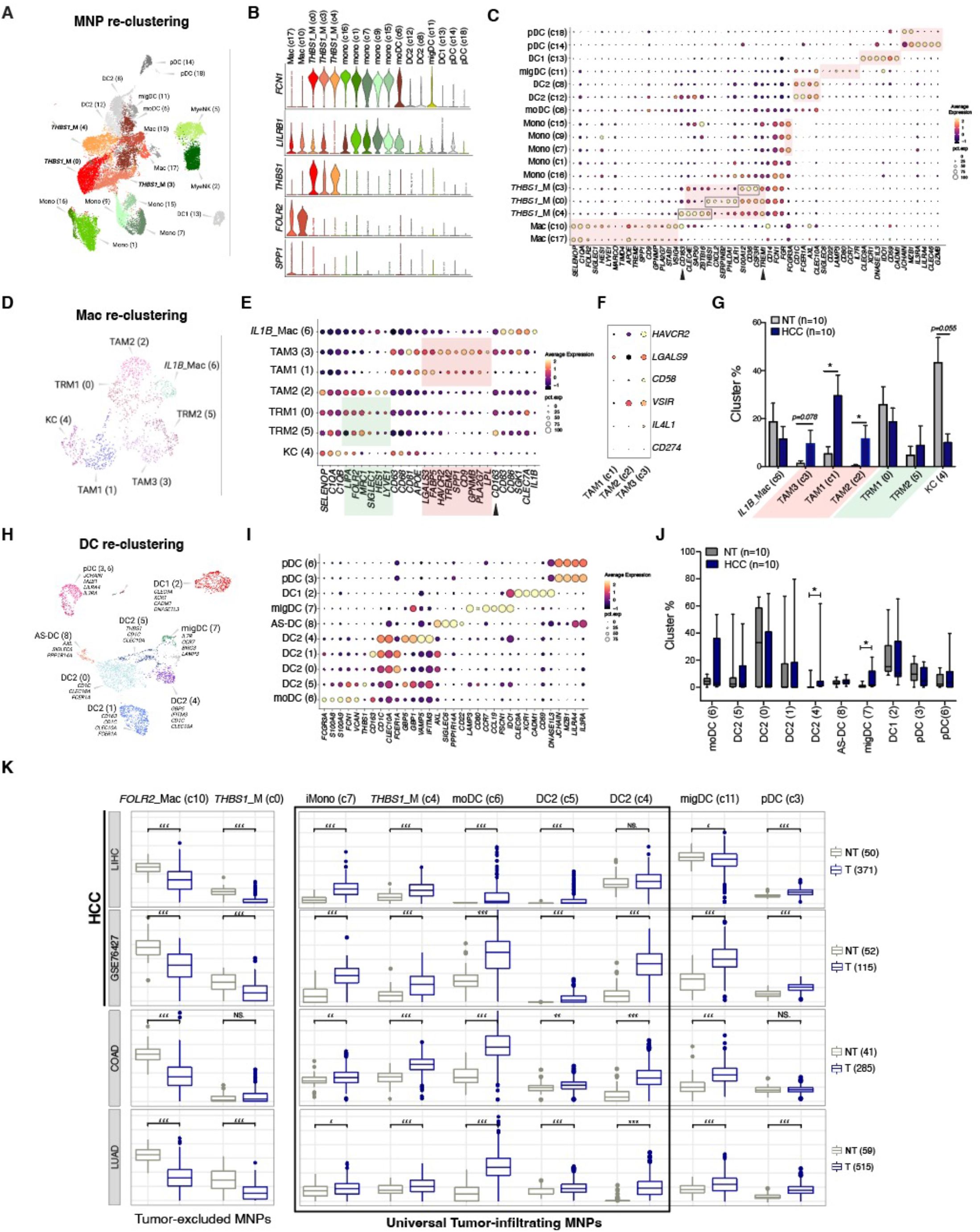
Defining MNP subsets and their predicted tumoral distribution. (A) Louvain clustering of 25,146 MNP transcriptome libraries colored based on cell type annotation. (B) Violin plots depicting the expression per cluster of selected discriminating genes (C) Dot plot depicting average expression (dot color) per cluster of selected discriminating genes and percentage of cells (dot size) expressing those genes. (D) Louvain re-clustering of macrophages (MNP clusters 10 and 17) identifies 7 new clusters of KC, TRMs, TAMs and an inflammatory *IL-1B*_Mac subset. (E) Dot plot depicting average expression per cluster of selected discriminating genes (F) Dot plot depicting average expression of immune checkpoint genes in TAMs. (G) Macrophage cluster frequencies in the adjacent NT (grey) versus HCC (blue) of the 10 patients investigated by scRNA-seq. *p<0.05 using multiple Wilcoxon tests (H) Louvain re-clustering of DCs (MNP clusters 8, 11, 12, 13, 14 and 18) (I) Dot plot depicting average expression per cluster of selected discriminating genes (J) DC cluster frequencies in the adjacent NT and HCC of the 10 patients investigated by scRNA-seq. *p<0.05 using multiple Wilcoxon tests (K) Box plots depicting the predicted proportions of MNP subsets by CIBERSORTx deconvolution of bulk transcriptomics datasets from four cohorts of patients, with either HCC (TCGA_LIHC; GSE76427), colon adenocarcinoma (TCGS_COAD) or lung adenocarcinoma (TCGA_LUAD). Signatures derived from the MNP re-clustering were used, except for the boxplots of DC2 (c4, c5) generated using DC re-clustering signatures.

### Specific populations of MNP infiltrate HCC primary tumors

To investigate MNP diversity and distribution in HCC, we re-clustered their transcriptomes. A total of 19 Louvain clusters were identified, including 17 clusters of MNPs and two clusters of MyeNK (c2, 5) (Figure 2a). The MNP clusters encompassed macrophages (c10, 17), a spectrum of monocyte-like states (c0, 1, 3, 4, 7, 9, 15, 16), monocyte-derived DC (moDC; c6) and DCs (c8, 11, 12, 13, 14, 18). *FCN1* expression marked moDCs and all monocyte states (Figure 2b). Among these, three clusters (c0, 3, 4), discriminated by the selective expression of *THBS1*, referred to as *THBS1_*Myeloid (*THBS1_*M), were enriched for signatures ascribed to myeloid-derived suppressor cells (MDSCs) (Condamine *et al*., 2016) (Figure 2b, c and Table S4). The other clusters (c1, 7, 9, 15, 16) expressed highest levels of *LILRB1, LILRB2* and *POU2F2* (Figure 2b, Table S4), features associated with differentiated monocytes (Weinreb et al., 2020). These could be classified according to *CD14* and *FCGR3A* (CD16) expression, into *CD14^+^CD16^-^* classical monocytes (cMono) (c1, 16), *CD14^+^CD16^+^* intermediate monocytes (iMono) (c7) and *CD14^-^CD16^+^* non-classical monocytes (ncMono) (c9, 15). A core signature (Table S5) of 6 genes (*SELENOP, APOE, C1QA, CD81, FOLR2, TREM2*) distinguished macrophages (c10, 17) from other innate immune cells (Figure S3a-d; Figure 2b, c). c10 expressed highest levels of *FOLR2*, previously described as a marker of tissue-resident macrophages (TRM) (Nalio Ramos et al., 2022), whereas c17 was discriminated by the expression of *SPP1* (Figure 2b), which marks tumor-associated macrophages (TAMs). However, c17 also expressed *FOLR2* at lower level, and Kupffer cell (KC) features e.g., *MARCO* and *TIMD4* (Figure 2c), suggesting mixed populations. To refine the macrophage characterization, we re-clustered c10 and c17, which distinguished seven macrophage subsets (Figure 2d): one inflammatory (c6; *IL1B, CD83, CD86*), two *FOLR2*^-^*TREM2*^+^ (c1, 3), expressing in common *CD9, SPP1* and *GPNMB* among others, and four *FOLR2^+^TREM2^-^* (c0, c2, c4, c5) (Figure 2d, e). The latter included KC (c4; *MARCO, TIMD4*) and three TRM-like clusters expressing higher levels of *MRC1 (CD206)* (Figure 2e and S4a). A previous study implicated NOTCH signaling in re-programming a subset of bone marrow (BM)-derived macrophages into *FOLR2^+^* TRM-like macrophages (Sharma *et al*., 2020). Cells in c2 expressed high levels of the NOTCH effector *HES1* (Figure 2e) and scored highly for our TRM signature (Figure S4b), suggesting that they might represent such a population. As reported in (Sharma *et al*., 2020), they highly expressed *LYVE1* (Figure 2e), which marks peri-vascular localization. Furthermore, our analysis revealed that among the *TREM2^+^SPP1^+^* cells, two subsets can be distinguished based on differential expression of *CD163,* a *CD163*^low^ subset (TAM1 (c1)) with discriminatory expression of metallothioneins (*MT1X, MT1G, MT2A*), and a *CD163*^high^ subset (TAM3 (c3)) expressing several immune checkpoints at higher levels compared to other TAM populations, including *HAVCR2* (TIM3), *LGALS9* (Galectin-9), *VSIR* (VISTA), *CD58* and the metabolic immune checkpoint interleukin-4-induced-1 (*IL4L1*) (Sadik et al., 2020) (Figure 2f). Notably, all three TAM subsets were enriched in HCC compared to NT in our patient cohort, while KCs were depleted (Figure 2g).

To explore the heterogeneity of DCs in HCC, we re-clustered the DC transcriptomes, which identified one DC1 cluster (c2) expressing *CLEC9A, XCR1* and *DNASE1L3*, one AS-DC (c8) expressing *AXL* and *SIGLEC6*, one migratory DC (migDC, c7) (Miller et al., 2012), also referred to as mature regulatory DCs (Maier et al., 2020), expressing maturation and migration genes *(LAMP3, CCL19, CCR7)*, and four DC2 states expressing in common *CD1C* and *CLEC10A*. *FCER1A* marked DC2 (c0, 1), with (c1) distinguished by high expression of *CD163*. c4 found in HCV-infected HCC patients exhibited a type I IFN signature *(IFITM3, GBP5, GBP1, VAMP5),* while c5 shared features with moDC, expressing monocyte-associated genes *(FCN1, S100A8/9),* and was marked by high *THBS1* expression. In addition, two clusters of pDC (c3, 6) were identified expressing *JCHAIN, LILRA4, MZB1,* and *IL3RA* (Figure 2h, i). Among the DC subsets, migDCs were significantly enriched in the tumor tissue of all patients in our cohort (Figure 2j), and DC2 (c4) found in HCV-infected patients were also enriched in HCC compared to NT (Figure 2j).

To confirm in large cohorts of patients the MNP subsets infiltrating HCC primary tumors, we next used the deconvolution algorithm CIBERSORTx (Steen et al., 2020), trained with our scRNA-seq data to predict cell-type density in bulk transcriptomics datasets. We first interrogated two cohorts of patients with HCC, i.e., TCGA_LIHC (371 patients) and GSE76427 (115 patients) (Grinchuk et al., 2018). We also explored MNP subset densities in two unrelated cohorts of patients with lung adenocarcinoma (TCGA_LUAD; 515 patients) and colon adenocarcinoma (TCGA_COAD; 285 patients). Our results show a consistent tumoral depletion of *FOLR2_*TRM (c10) in all cohorts (Figure 2k), in line with findings in human breast cancer (Nalio Ramos *et al*., 2022). In contrast, iMono (c7) and moDC (c6) are more abundant in the tumor compared to NT in all cohorts. With the exception of LIHC, migDC (c11) infiltrate the tumor in most cohorts, i.e., LUAD, COAD and GSE76427, consistent with our scRNA-seq results (Figure 2k). cMono (c1) and ncMono (c9) are inversely correlated in HCC, with lower cMono (c1) proportions in the tumors. However, this tissue distribution is not consistent in LUAD and COAD (Figure S4c), suggesting HCC-specific regulation. Surprisingly, the three MDSC-like *THBS1_*M subsets, do not display similar patterns of tumoral infiltration. Whereas *THBS1_*M (c0) are depleted from the tumor in HCC and LUAD, *THBS1_*M (c4) significantly infiltrate the tumor along with moDC (c6) and iMono (c7) in all cohorts (Figure 2k). The densities of *THBS1_*M (c3) and *SPP1_*Mac (c17) is variable among tissues and across cohorts (Figure S4c). Similar results were obtained when we deconvoluted using subset expression profiles from the macrophage and DC re-clustering. This analysis further unraveled a pDC subset (c3) that infiltrate tumors in HCC and LUAD, and two inflammatory DC2 subsets (c4, c5) as additional tumor-infiltrating MNPs in all cohorts (Figure 2k).

Since HCC etiology might underlie the heterogeneity of response to ICB in patients with HCC (Pfister *et al*., 2021), we took advantage of our scRNA-seq data to assess the impact of etiology on myeloid cell diversity in HCC tumors. Our results show that among the tumor-infiltrating MNPs, *THBS1_*M (c4) cluster was exclusively found in patients with steatohepatitis but not in viral HCC (Figure S4d, e). ncMono (c9, 15) and DC2 (c12) also expanded in this condition (Figure S4d, e). In contrast, iMono (c7) and moDC (c6) were less abundant in steatohepatitis compared to viral HCC (Figure S4d, e). Collectively, our results identify iMono, moDC, inflammatory DC2 and the MDSC-like *THBS1_*M (c4) subset, as universal tumor-infiltrating MNPs (TIMs) and unravel an effect of steatohepatitis on intra-tumoral MNP states, particularly an expansion of the MDSC-like *THBS1_*M (c4) population.

### Fine-mapping MDSC-like states in HCC

Despite their significance in cancer progression, MDSCs are poorly annotated, and the phenotypic markers used to isolate them overlap with those of normal neutrophils and monocytes in both mice and humans (Veglia et al., 2021). Our analysis revealed three clusters of *THBS1*_M cells enriched in MDSC signatures (Condamine *et al*., 2016). These outnumbered TAMs in HCC, representing 27.4% of all MNP analyzed (c0, 10.8%; c3, 8.4%; c4, 8.2%) (Figure 2a). To better characterize them, we first elucidated a core signature of 10 genes (*THBS1, TREM1, S100A12, SERPINB2, OLR1, SAP30, PHLDA1, MEGF9, VEGFA, VCAN*) that identified them primarily in c10 (and c5 to a lower extent) in the innate immune dataset (Figure 3a, b), and distinguished them from other MNP (Figure S3m, n). They were selectively marked by *THBS1,* and expressed both monocyte-lineage (*FCN1*) and neutrophil-lineage (*CSF3R*) genes, but were not fully differentiated monocytes (*POU2F2*) (Weinreb *et al*., 2020)*. TREM1* encoding ‘Triggering receptor expressed on myeloid cells 1’, highly expressed in neutrophils (Figure 3c), discriminated the three *THBS1_*M clusters from other MNPs (Figure 3d). The lectin type oxidized LDL (oxLDL) receptor 1 (LOX-1, encoded by *OLR1*), previously described as a specific marker of human PMN-MDSC (Condamine *et al*., 2016), marked c0 (Figure S5a-c). However, it was also detected in other monocytes at lower levels (Figure S5b, c), as reported in (Travaglini et al., 2020). In contrast, we identified *SERPINB2* that encodes the plasminogen activator inhibitor type 2 as a more robust marker of *THBS1_*M (c0) (Figure S5a-c). *MEGF9* (Multiple EGF Like Domains 9) was most highly expressed in *THBS1_*M (c3), while SAP30 (Sin3A Associated Protein 30) discriminated *THBS1_*M (c4) (Figure S5a, b). In order to elucidate the nature of the three *THBS1_*M subsets, we next performed an analysis of differentially expressed genes (DEG). This revealed that genes highly expressed in neutrophils e.g. *CXCL8, NAMPT* and *PLAUR* were also highly expressed in c0 as compared to c3/c4 (Figure 3e and Table S6), suggesting that c0 might encompass cells with a PMN-MDSC-like state. c3 was discriminated from c4 by monocyte lineage genes *ATF3, CD14, CSF3R, CACYBP, LGALS2, LILRA5* (Figure 3f and Table S6), suggesting that it may represent an M-MDSC-like state. In contrast, c4 had multi-lineage gene expression; its top DEGs encompassed macrophage- and DC-lineage genes (*C1QA/B/C, CD163* and *CLEC10A, CST3*), in addition to genes shared with c0 (*NAMPT, OLR1, PHLDA1/2, PLAUR*) (Figure 3f, g and Table S6). To explore the dynamic regulation of the three *THBS1_*M subsets, we next mapped their position along an inferred trajectory through Pseudotime ordering, which revealed three main states (Figure 3h). Analysis of temporally expressed genes along the Pseudotime identified the different profiles including a state expressing a variety of myeloid lineage genes at intermediate levels (Figure 3i). This subset was marked by high expression of inflammatory cytokine receptor genes e.g. *IL1RN, IL6R* and the inflammasome gene *NLRP3* (Figure 3i). Calculation of differentiation potency estimates revealed that the ‘multi-lineage’ *THBS1_*M (c4) had higher entropy rates compared to *THBS1_*M (c0), *THBS1_*M (c3) and *SPP1_*Mac (c17) (Figure 3j). Monocle2 pseudotime trajectory analysis revealed that *THBS1_*M (c4), moDC (c6) and iMono (c7) occupied an intermediate state along the pseudotime (Figure S5d). Similar results were obtained on Principal Component analysis of all MNP subsets, which segregated monocytes, from macrophages and DCs, and revealed intermediate states for *THBS1_*M (c4), moDC (c6), iMono (c7) as well as migDC (c11) (Figure S5e). Predicted ordering by CYTOTRACE further supported the intermediate state of these subsets between DCs and macrophages/classical monocytes (Figure S5f). Unlike *THBS1_*M (c4), *THBS1_*M (c0) and *THBS1_*M (c3) exhibited a more committed state and segregated closer to differentiated monocytes (Figure S5e, f). Together, our results unravel a discrete population of MDSC-like cells (*THBS1_*M (c4)) that consistently infiltrate tumors and are distinguished from other MNPs by ‘multi-lineage’ expression of granulocyte and macrophage/DC genes.

**Figure 3.**
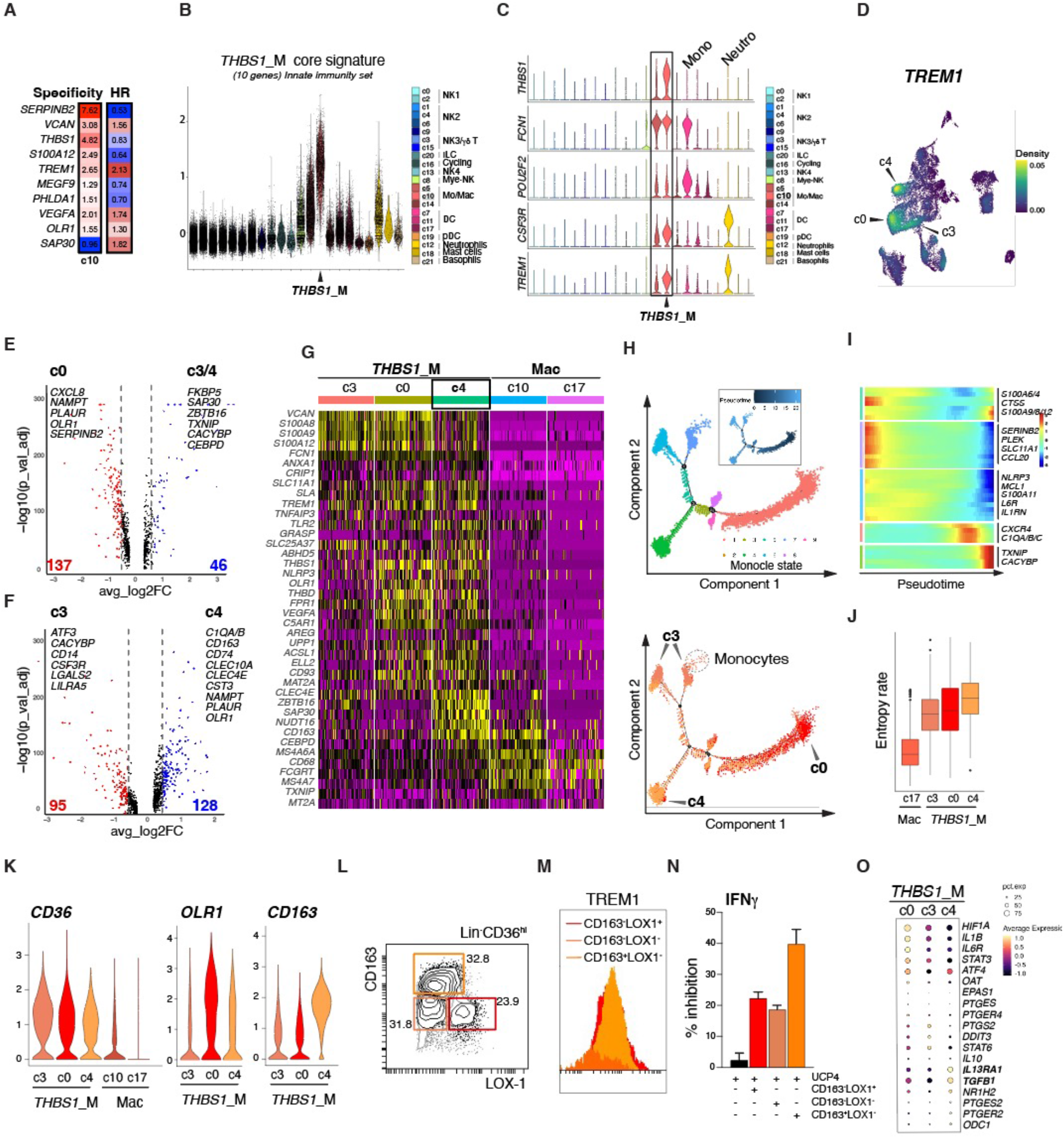
Identification of *TREM1^+^CD163^+^* M_reg_ cells as potent immunosuppressive myeloid cells. (A) *THBS1_*M (c10) core signature specificity and association with mOS in patients with HCC from TCGA_LIHC cohort. HR, hazard ratio (B) Violin plots depicting the signature score of *THBS1_*M 10 genes-core signature in the different innate immune main clusters. (C) Violin plots depicting the expression per cluster of selected discriminating genes in the different innate immune main clusters. (D) Density plot representing the expression of *TREM1* as a discriminatory marker of the three *THBS1_*M clusters. (E) Volcano plot depicting differentially expressed genes between *THBS1_*M (c0) and other *THBS1_*M cells (c3+c4). (F) Volcano plot depicting differentially expressed genes between *THBS1_*M (c3) and *THBS1_*M (c4). (G) Heatmap representing expression of selected genes from CIBERSORTx top 60 genes in (c4) for 100 cells in each cluster shown. (H) Pseudotime trajectory analysis using Monocle2 of the three THBS1_M clusters, colored by monocle state, Pseudotime or cluster. (I) Heatmap of the top temporally expressed genes identifying the different Monocle2 states. (J) Box plots of signaling entropy values (y-axis, also named potency) in the three *THBS1*_M clusters and the *SPP1*_Mac cluster. (K) Violin plots depicting expression of *CD36, OLR1* and *CD163* in the indicated MNP clusters. (L) Gating strategy used to sort human *THBS1_*M (c0, c3 and c4 of the MNP dataset) subsets according to differential expression of LOX-1 and CD163 in the Lin^-^CD36^+^ population. M) Histograms depicting TREM1 expression among the different *THBS1_*M subsets according to differential expression of LOX-1 and CD163 in the Lin^-^CD36^+^ population. (N) Quantification of the % inhibition of IFNγ production by a TERT-specific CD4^+^ T cell clone stimulated or not with UCP4 TERT peptide in co-culture with different *THBS1_*M subsets sorted from HCC surgical resections according to the gating strategy in (L). (O) Dot plot depicting average expression per cluster of genes associated with myeloid immunosuppressive functions.

### TREM1^+^ CD163^+^ myeloid cells are super immunosuppressors

We next sought to investigate the function of the three *THBS1_*M populations. We searched for surface proteins differentially expressed among them. We found that the scavenger receptor CD36 was highly expressed on all three *THBS1_*M subsets compared to macrophages (Figure 3k) and other MNPs (Figure 2c). Among the *THBS1_*M subsets, *OLR1* was mainly expressed in *THBS1_*M (c0) with intermediate levels in *THBS1_*M (c4), whereas CD163 was highly expressed in *THBS1_*M (c4) (Figure 3k). Flow cytometry analysis of CD163 and LOX-1 revealed mutually exclusive staining within the CD45^+^CD3^-^CD19^-^CD56^-^CD36^+^ population (Figure 3l). Consistent with our scRNA-seq data, surface expression of TREM1 was more prominent on the LOX-1^+^CD163^-^ (*THBS1_*M (c0)) and LOX-1^-^CD163^+^ (*THBS1_*M (c4)) populations, as compared to LOX-1^-^CD163^-^ (*THBS1_*M (c3)) cells (Figure 3m). The *THBS1_*M (c4) subset can thus be identified as the main immune population in HCC dually expressing TREM1 and CD163. Because of their high entropy and enrichment in MDSC signature genes, we next functionally assessed their immunosuppressive activity. We FACS-sorted the three subsets from surgical resections of patients with HCC and co-cultured them with a high avidity Th1-polarized CD4^+^ T cell clone reactive to the tumor-associated antigen (TAA) Telomerase (TERT) peptide UCP4, as in (Lauret Marie Joseph et al., 2020). All three subsets inhibited the production of IFNγ by CD4^+^ T cells upon UCP4 stimulation, but the TREM1^+^CD163^+^ population showed the most potent inhibitory effect (Figure 3n and S5g). As a control, we generated human monocyte-derived suppressor cells (HuMoSc) *ex vivo* according to an established protocol (Janikashvili et al., 2015), and confirmed that they were highly efficient in suppressing cytokine production by activated T cells (Figure S5h, i). Interestingly, phenotypic characterization of these HuMoSc showed that they were primarily CD163^+^ (>85%) and highly expressed TREM1 on their surface (Figure S5j, k), suggesting that a strong immunosuppressive capacity can be identified by dual expression of TREM1 and CD163 on monocytic cells. Myeloid cells exert their immunosuppressive functions through different mechanisms. e.g. via reactive oxygen species (ROS), nitric oxide (NO), arginase, prostaglandin E2 (PGE_2_), anti-inflammatory cytokines (e.g. IL-10, TGFβ) and cell surface immune checkpoints such as PD-L1 (Hegde et al., 2021). Interrogation of the expression of central effectors in these pathways revealed that *TREM1^+^CD163^+^* myeloid cells (*THBS1_*M (c4)) expressed highest levels of *TGFB1* (Figure 3o), which was also observed in *ex vivo* generated HuMoSc (Figure S5l). They also exhibited highest levels of *IL13RA*, which encodes a chain of the receptor for IL-13, a cytokine implicated in TGFβ induction (Fichtner-Feigl et al., 2006; Lee et al., 2001). In addition, they had elevated levels of *PTGES2* and *PTGER2,* encoding a PGE2 synthase and receptor, respectively, as well as *ODC1,* encoding ornithine decarboxylase that acts downstream of arginase as the rate-limiting enzyme in the polyamine biosynthesis pathway. In addition, they shared with *THBS1_*M (c0) high expression of *ATF4* (encoding Activating Transcription Factor 4, an endoplasmic reticulum (ER) stress effector recently demonstrated to be required for MDSC immunosuppressive function (Halaby et al., 2019)) (Figure 3o). Collectively, these results uncover a discrete population of regulatory myeloid cells (M_reg_), identified in HCC as TREM1^+^CD163^+^, with potent immunosuppressive function.

### *TREM1^+^CD163^+^* M_reg_ cells are enriched at fibrotic lesions in HCC

Liver fibrosis is a dominant precursor of HCC irrespective of etiology and the IL-13-TGFβ1 axis is a major driver of this process (Chiaramonte et al., 1999; Nakamura et al., 2000; Roulot et al., 1999). TGFβ activates hepatic stellate cell (HSC), promotes their function in extracellular matrix remodeling and collagen deposition (Hellerbrand et al., 1999), and mediates resistance to anti-PD-L1 treatment (Mariathasan et al., 2018). IL-13 receptor signaling elicits TGFβ production (Fichtner-Feigl *et al*., 2006; Lee *et al*., 2001), but also cooperates with it in independently inducing the expression of pro-fibrotic genes such as matrix metalloproteases (MMPs), tissue inhibitors of metalloprotease-1 (TIMP1) and interstitial collagens (Gieseck et al., 2016; Kaviratne et al., 2004), and IL13RA blockade has been shown to inhibit hepatic fibrosis (Chiaramonte *et al*., 1999). Our results revealed highest expression of *IL13RA* and *TGFB1* in *TREM1^+^CD163^+^*myeloid cells (*THBS1_*M (c4)). We therefore wondered whether these cells were associated with HCC fibrotic lesions. To this end, we performed spatial transcriptomics (ST) on tumor tissue sections from patients with HCC. Two patients had confirmed fibrosis: patient #20 had severe fibrosis (Metavir, F3) linked to alcohol consumption, while patient #23 had moderate fibrosis (Metavir, F1/F2) linked to NASH (Table S1). Fibrotic lesions were evident on the H&E-stained tissue sections processed for ST (Figure 4a, f). Transcriptomics from 2261 and 1987 spots were obtained at a median depth of 34,526 and 26,017 UMIs/spot, and 5882 and 4954 genes/spot for patients # 20 and #23, respectively (Table S7). Unbiased clustering identified eight clusters in patient #20 and ten clusters in patient #23 (Figure 4b, g). Based on spot feature expression of a fibrosis-associated signature (*COL1A1*, *TIMP1, ACTA2*), we identified clusters 2 and 5 in patient #20, and cluster 8 in patient #23 as scoring highly for the fibrosis signature (Figure 4c-d, h-i). We next interrogated the *in situ* expression of a geneset composed of four genes (*TREM1, VCAN, THBS1, S100A4*) collectively discriminating *THBS1_*M cells from other innate immune cells (Figure 4k). Our results revealed a striking overlap in the expression of this gene signature with that marking fibrotic lesions (Figure 4d-e, i-j). *VCAN, THBS1* and *S100A4* may also be expressed by fibroblasts indicating that *THBS1_*M cells are geared with a pro-fibrotic machinery, consistent with their high expression of *TGFB1* (Figure 3o) and *TIMP1* (Figure 4k). To ensure specific detection of *TREM1^+^CD163^+^*M_reg_ cells, we next examined the expression of a more-restricted signature composed of *TREM1, CD163* and *CLEC4E* that discriminates this subset from other MNPs (Figure 2c and Table S4). Our results revealed that *TREM1^+^CD163^+^* M_reg_ cells populate fibrotic lesions (Figure 4d-e, i-j).

**Figure 4.**
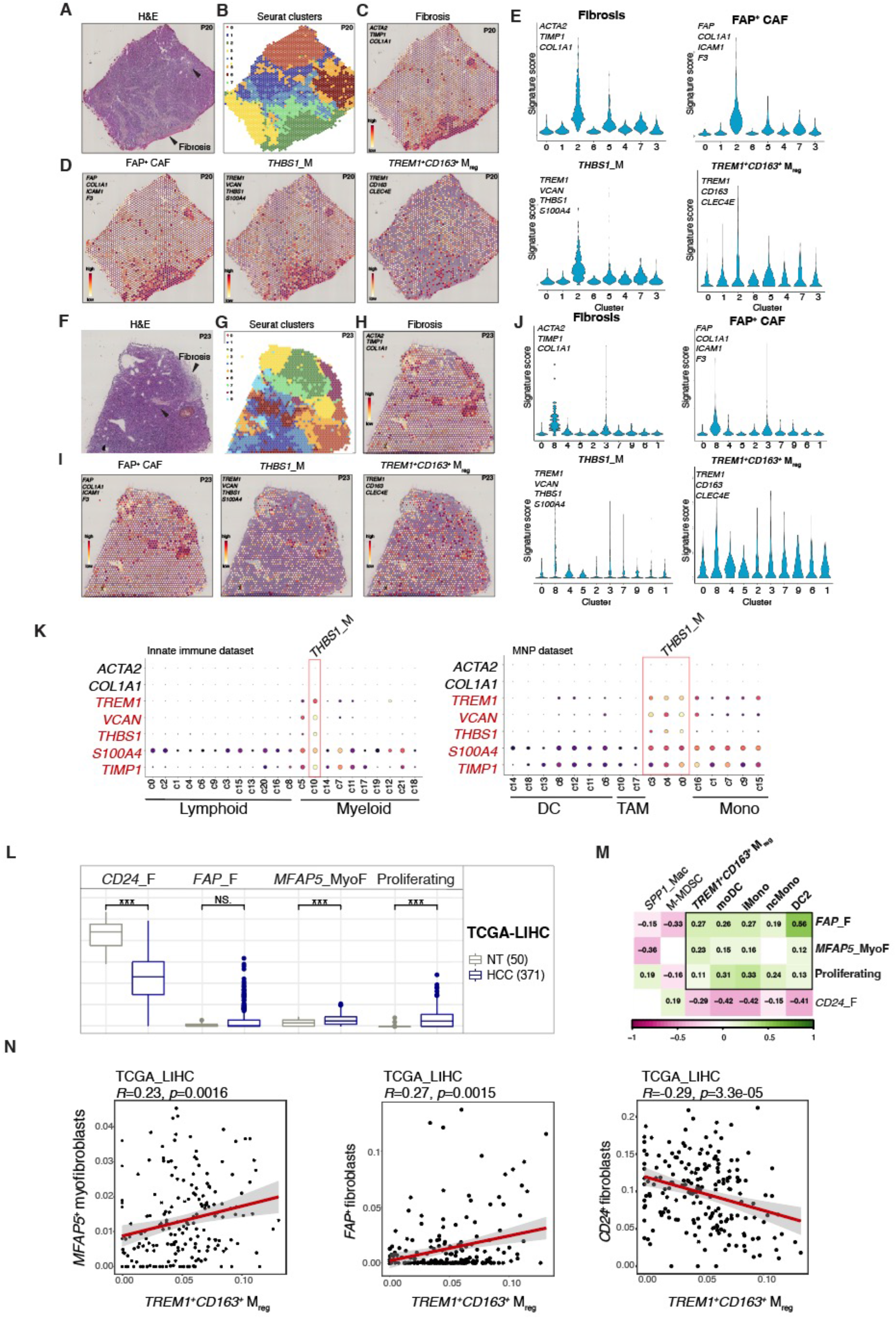
*TREM1^+^CD163^+^* M_reg_ cells populate fibrotic lesions in HCC. (A-J) Tumor tissue sections from two patients with HCC (P20, P23) were analyzed by stRNA- seq using the Visium platform from 10x Genomics. (A, F) Tumor tissue sections stained with H&E (B, G) Overlapping spatial localization of spatial Louvain Seurat clusters (C, H) Spatial scoring of a gene signature marking fibrosis (*ACTA2, TIMP1, COL1A1*) (D, I) Spatial scoring of gene signatures identifying FAP^+^ cancer-associated fibroblasts (CAFs) (*FAP, COL1A1, ICAM1, F3*) (left), *THBS1*_M (*TREM1, VCAN, THBS1, S100A4*) (middle) and *TREM1^+^CD163^+^* M_reg_ (*TREM1, CD163, CLEC4E*) (right). (E, J) Violin plots depicting signature score of the interrogated spatial genesets in each Seurat cluster. (K) Dot plots depicting average expression per cluster of genes associated with fibrosis or expressed in *THBS1*_M in the innate immunity dataset (left) or MNP dataset (right). (L) Box plots depicting the predicted proportions of fibroblast subsets by CIBERSORTx deconvolution of bulk transcriptomics data from the TCGA_LIHC cohort. (M) Matrix of Pearson correlations from predicted proportions of MNP and fibroblast subsets deconvoluted from the TCGA_LIHC cohort data. Green, positive correlation; Pink, negative correlation (N) Scatter plots depicting Pearson correlations between proportions of *TREM1^+^CD163^+^* M_reg_ and different fibroblast subsets.

The heterogeneity of fibroblasts shapes the tumor immune microenvironment and is associated with the outcome of cancer progression (Davidson et al., 2021; Peltier et al., 2022). Specific cancer-associated fibroblast (CAF) sub-populations promote immunosuppression in the TME and have been linked to immunotherapy resistance (Kieffer et al., 2020; Qi et al., 2022). Immunosuppressive CAFs exhibit an activated phenotype and express markers such as fibroblast-activation protein (*FAP*) and components of the desmoplastic structures, including collagens (*COL1A1, COL3A1*), integrins (*ICAM1*) and tissue factor (*F3*). We interrogated the spot feature expression of a FAP^+^ CAF signature (*FAP, COL1A1*, *ICAM1, F3*) in our ST data and observed overlapping spatial distribution with that of *TREM1^+^CD163^+^* M_reg_ (Figure 4d-e, i-j), suggesting potential crosstalk between these two populations. To validate this observation in a larger cohort of HCC patients, we used CIBERSORTx deconvolution to predict the abundance of MNP subsets defined in our study and that of 7 CAF populations defined by scRNA-seq in (Qi *et al*., 2022), namely *FAP*_fibroblasts, MFAP5_myofibroblasts, proliferating fibroblasts, *CD24_*fibroblasts, *FGFR2_*fibroblasts, *CD73_*fibroblasts and DES_fibroblasts. Our results show that *FAP*_fibroblasts, *MFAP5*_myofibroblasts and proliferating fibroblasts were enriched in the tumors of HCC patients, whereas *CD24_*fibroblasts were not (Figure 4l). We next performed Pearson correlations to assess infiltration patterns of MNP and CAF subsets. We observed significant positive correlations between *TREM1^+^CD163^+^*M_reg_ and *FAP*_fibroblasts, *MFAP5*_myofibroblasts and proliferating fibroblasts (Figure 4m, n). The same correlations were observed for iMono (c7), ncMono (c9), moDC (c6) and DC2 (c8). Together, these results point to a specific spatial co-localization of *TREM1^+^CD163^+^* M_reg_ with pro-fibrogenic, immunosuppressive CAFs in tumors from patients with HCC.

### *TREM1^+^CD163^+^* M_reg_ correlate with poor clinical outcomes and reduced survival in patients with HCC and predict resistance to immune checkpoint blockade

Suppressive myeloid cells are collectively considered as tumor promoting and an impediment to efficient anti-tumor immunity. Our data revealed significant tumor infiltration of *TREM1^+^CD163^+^*M_reg_ in several cohorts of patients with solid tumors, including HCC (Figure 2k). TREM1^+^CD163^+^ M_reg_ have superior immunosuppressive activity (Figure 3m) and specifically associate with tumor-promoting and immunosuppressive *FAP^+^* CAFs (Figure 4m). Therefore, we wondered whether *TREM1^+^CD163^+^*M_reg_ are associated with worse survival in patients with HCC. We first explored the median expression of the *THBS1_*M core signature geneset (Table S5) in the TCGA_LIHC cohort. Patients with higher expression of this geneset have significantly poorer mean overall survival (mOS) (Figure 5a). We next used CIBERSORTx deconvolution to interrogate MNP associations with tumor grade. PMN-MDSC-like *THBS1_*M (c0), M-MDSC-like *THBS1_*M (c3), *FOLR2*_Mac (c10), and *SPP1*_Mac (c17) are not associated with this parameter (Figure S6a). In contrast, *TREM1^+^CD163^+^* M_reg_ significantly associate with higher pathological tumor grade (stages III and IV) (Figure 5b), consistent with our scRNA-seq data in which cells of this subset were exclusively derived from patients with high grade HCC (patients #20 and #23; Table S2). iMono (c7) and moDC (c6) are also enriched in higher grade tumors (Figure 5b). Second, we assessed links with serum alpha-fetoprotein (AFP) levels, an important prognostic factor and a predictive biomarker of mortality (Tsukuma et al., 1993). Among the *THBS1_*M populations, only the proportion of *TREM1^+^CD163^+^* M_reg_ strongly correlates with elevated AFP (Figure 5c), whereas M-MDSC-like *THBS1_*M (c3) are counter-correlated (Figure S6b). iMono and moDC have significant albeit weaker associations with this clinical parameter (Figure 5c). Third, we examined associations with the inactivation of HIPPO, as defined by the silence of HIPPO [SOH] signature, reported to predict poor prognosis in patients with HCC (Sohn et al., 2016). The proportions of *TREM1^+^CD163^+^*M_reg_ and moDC but not that of other MNPs are higher in the SOH class of patients compared to patients with active HIPPO signaling (AH) (Figure 5d and S6c). Last, we interrogated an integrative classification by Hoshida *et al*. that stratifies patients according to molecular and clinical features (Hoshida et al., 2009). As for pathologic tumor grade, serum AFP and SOH, the proportion of *TREM1^+^CD163^+^*M_reg_ are highest in the Hoshida S2 subclass, characterized by larger and poorly differentiated tumors, elevated serum AFP and poor mOS (Figure S6d). M-MDSC-like *THBS1_*M (c3) do not associate with the Hoshida classification, whereas the proportions of PMN-MDSC-like *THBS1_*M (c0) and *FOLR2_*Mac decrease with HCC severity (S3→S1→S2) (Figure S6d). Collectively, these results identify *TREM1^+^CD163^+^* M_reg_, moDC and iMono as the main tumor-infiltrating MNPs that associate with poor prognosis in patients with HCC, according to the measured clinical parameters.

**Figure 5.**
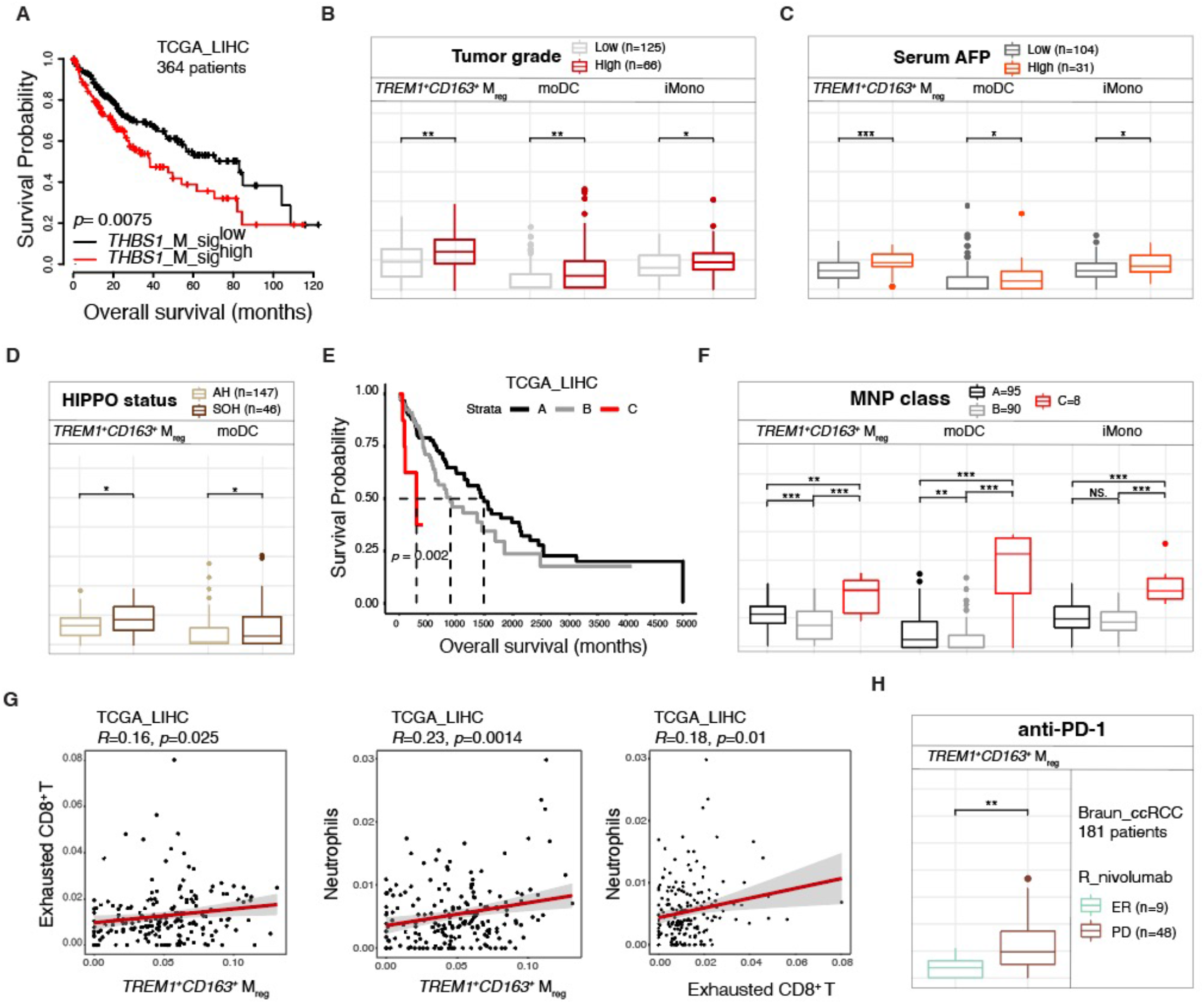
*TREM1^+^CD163^+^*M_reg_ correlate with poor prognosis in HCC and resistance to immune checkpoint blockade. (A) Kaplan-Meier survival curves of patients with HCC (n=364) (Menyhart et al., 2018) according to expression of the *THBS1_*M core signature. (B-D) Box plots depicting the predicted proportions of MNP subsets by CIBERSORTx deconvolution of bulk transcriptomics data from the TCGA_LIHC cohort according to tumor grade (Low, pathologic T grade 1 and 2; High, pathologic T grade 3 and 4) (B), serum AFP (<300 or >300) (C), or HIPPO status (D). (E) Kaplan-Meier survival curves of patients with HCC (n=193) from TCGA_LIHC cohort stratified according to MNP infiltrate profiles. (F) Box plots depicting the predicted proportions of MNP subsets according to patient group from (E). (G) Scatter plots depicting Pearson correlations between proportions of *TREM1^+^CD163^+^* M_reg_ and exhausted CD8^+^ T cells or neutrophils, and Pearson correlations between proportions of neutrophils and exhausted CD8^+^ T cells. (H) Box plots depicting the predicted proportions of *TREM1^+^CD163^+^* M_reg_ according to patient response to anti-PD-1 (Nivolumab) in a cohort of patients with clear cell renal cell carcinoma (ccRCC).

Next, we wondered whether the nature of the myeloid infiltrate could stratify HCC patients and serve as an immunological biomarker of clinical outcomes. To address this, we performed hierarchical unsupervised clustering of HCC patients in the TCGA_LIHC cohort based on CIBERSORTx predicted proportions of the MNP populations we characterized in this study. Tumor-infiltrating MNP profiles divided HCC patients into three strata (Figure S6e) with significantly different mOS (Figure 5e). Group A patients with the best mOS have higher proportions of *FOLR2*_Mac and *THBS1_*M (c0) (Figure S6e, f), i.e., the two MNP subsets generally depleted from solid tumors (Figure 2k), whereas higher *SPP1_*Mac proportions distinguishes group B patients with an intermediate mOS (Figure S6e, f). Group C patients, with markedly reduced mOS, are characterized by the highest proportions of *TREM1^+^CD163^+^* M_reg_ (*THBS1_*M (c4)), moDC and iMono (Figure 5f and S6e, f). Notably, the proportions of *TREM1^+^CD163^+^* M_reg_ in this cohort correlate with regulatory T cells (T_reg_), exhausted CD8^+^ T cells and neutrophils, according to expression profiles defined by scRNA-seq in (Qi *et al*., 2022) (Figure 5g). Last, we wished to examine myeloid subsets association with resistance to immune checkpoint blockade (ICB). Because we did not have access to transcriptomic data from study cohorts of patients with HCC treated with ICB, we interrogated datasets from other solid tumors. In a cohort of 181 patients with advanced clear cell renal cell carcinoma (ccRCC) treated with anti-PD-1 in prospective clinical trials (Braun et al., 2020), *TREM1^+^CD163^+^* M_reg_ density significantly associates with progressive disease (PD) (Figure 5h). None of the 18 other MNP cell subsets defined in this study correlate with immunotherapy resistance in this cohort (Figure S6g). Together, our results identify *TREM1^+^CD163^+^* M_reg_ as a potent immunosuppressive tumor-infiltrating myeloid population associated with poor prognosis in HCC and resistance to ICB described in ccRCC.

### TREM1 associates with poor prognosis in solid tumors and confers immunosuppression

Among the 10 genes constituting the *THBS1_*M core signature, *TREM1* confers the highest hazard ratio on HCC survival in the TCGA_LIHC cohort (Figure 3a). Patients with high *TREM1* expression have significantly poorer mOS than patients with lower expression (Figure 6a), consistent with previous reports in other HCC patient cohorts (Duan et al., 2015; Liao et al., 2012). Our findings that *TREM1^+^CD163^+^* M_reg_ are highly immunosuppressive and predict poor prognosis in HCC prompted us to interrogate whether *TREM1* expression *per se* is associated with poor clinical outcomes, and if it plays a direct role in immunosuppression. We first interrogated *TREM1* expression in the three HCC patient groups from the TCGA_LIHC cohort classified according to MNP infiltration profiles. We observed highest expression of *TREM1* in group C patients (Figure 6b), enriched in *TREM1^+^CD163^+^* M_reg_ and with a poor mOS (Figure 5e, f). In addition, *TREM1* expression is higher in patients with fibrosis, and significantly associates with the silence of HIPPO (SOH) and the Hoshida S2 class (Figure 6b). *TREM1* positively correlates with several effectors of immunosuppression, most significantly with *TGFB*, *IL10, PTGES2* and *IL8* (Figure 6f), as well as with *PTGER2* and *CD274* (PD-L1) (Figure S7a). Besides HCC, *TREM1* expression also associates with high pathologic tumor grades in other solid tumors, namely colon and rectum adenocarcinoma and kidney renal clear cell carcinoma, as revealed in TCGA_COAD, TCGA_READ and TCGA_KIRC cohort analyses, respectively (Figure S7b). Furthermore, *TREM1* expression significantly associates with resistance to anti-PD-1 revealed in the GSE78220 cohort of patients with melanoma (Hugo et al., 2017) (Figure S7c). Together, these results point to TREM1 as a potential effector of tumor progression and immunosuppression. To experimentally address the role of TREM1 in cancer immunity, we investigated its expression and function in mouse models of liver injury (Farrell et al., 2019), NASH and HCC *in vivo*, as well as in human monocyte cultures *ex vivo.* We used obese *db/db* diabetic mice fed a methionine and choline-deficient diet (MCD) for 7 weeks to elicit NASH, and carbon tetrachloride (CCL4)-treatment of C57Bl/6J mice as a model of liver injury (Figure S7d, e). In both disease models, mice had elevated levels of circulating liver enzymes, namely aspartate aminotransferase (AST) and alanine aminotransferase (ALT), indicative of liver damage (Figure S7f). In the NASH model, mice had elevated triglycerides, as expected, compared to control or CCL4-treated mice (Figure S7f). Immunophenotyping of the liver revealed enhanced accumulation of TNFα-producing CD8^+^ T cells in both NASH- affected or CCL4-treated mice (Figure S7g). In contrast, a significant influx of TREM1^+^ CD11b^+^ Ly6C^+^ Ly6G^-^ myeloid cells was observed in the NASH setting but not in control mice and to a lesser extent in CCL4-treated mice (Figure S7h), accompanied by an induction of TREM1 surface expression (Figure S7i), consistent with our scRNA-seq results revealing the *TREM1^+^CD163^+^* M_reg_ subsets in steatohepatitis-affected patients with HCC (Figure S4d, e). To address the role of TREM1 in HCC, we next used the Hep55.1C liver orthotopic mouse model of HCC and inhibited TREM1 signaling using the GF9 peptide (Shen and Sigalov, 2017) that interferes with TREM1 binding to its signaling adaptor DAP12. Our results show that pan- blockade of TREM1 signaling promoted tumor eradication (Figure S7j), as observed by others in different animal models (Wu et al., 2012; Wu et al., 2019). However, since the GF9 peptide could modulate TREM1 functions in immune and non-immune compartments, we next used the *ex vivo* HuMoSc model in co-culture with T cells to formally examine the role of TREM1 in the immunosuppressive functions of myeloid cells. Treatment of HuMoSc cells with the TREM1 ligand PGLYRP1 (peptidoglycan recognition protein 1) in complex with peptidoglycan (PGN) (PP) significantly increased cell surface expression of TREM1 (Figure 6g), and induced LOX-1 and CD15 (Figure 6h). In addition, it resulted in heightened production of IL-10 and IL-8 (Figure 6i), and processing and release of soluble TREM1 (sTREM1) (Figure 6i). Importantly, TREM1 stimulation led to potent suppression of T cell proliferation (Figure 6j) and effector functions, e.g. inhibition of TNFα production (Figure 6k). Together, these results suggest that the steatohepatitis-environment promotes the genesis of TREM1^+^ M_reg_ cells with strong immunosuppressive and pro-tumoral functions, and point to *TREM1* as a potential immunotherapeutic target in HCC.

**Figure 6.**
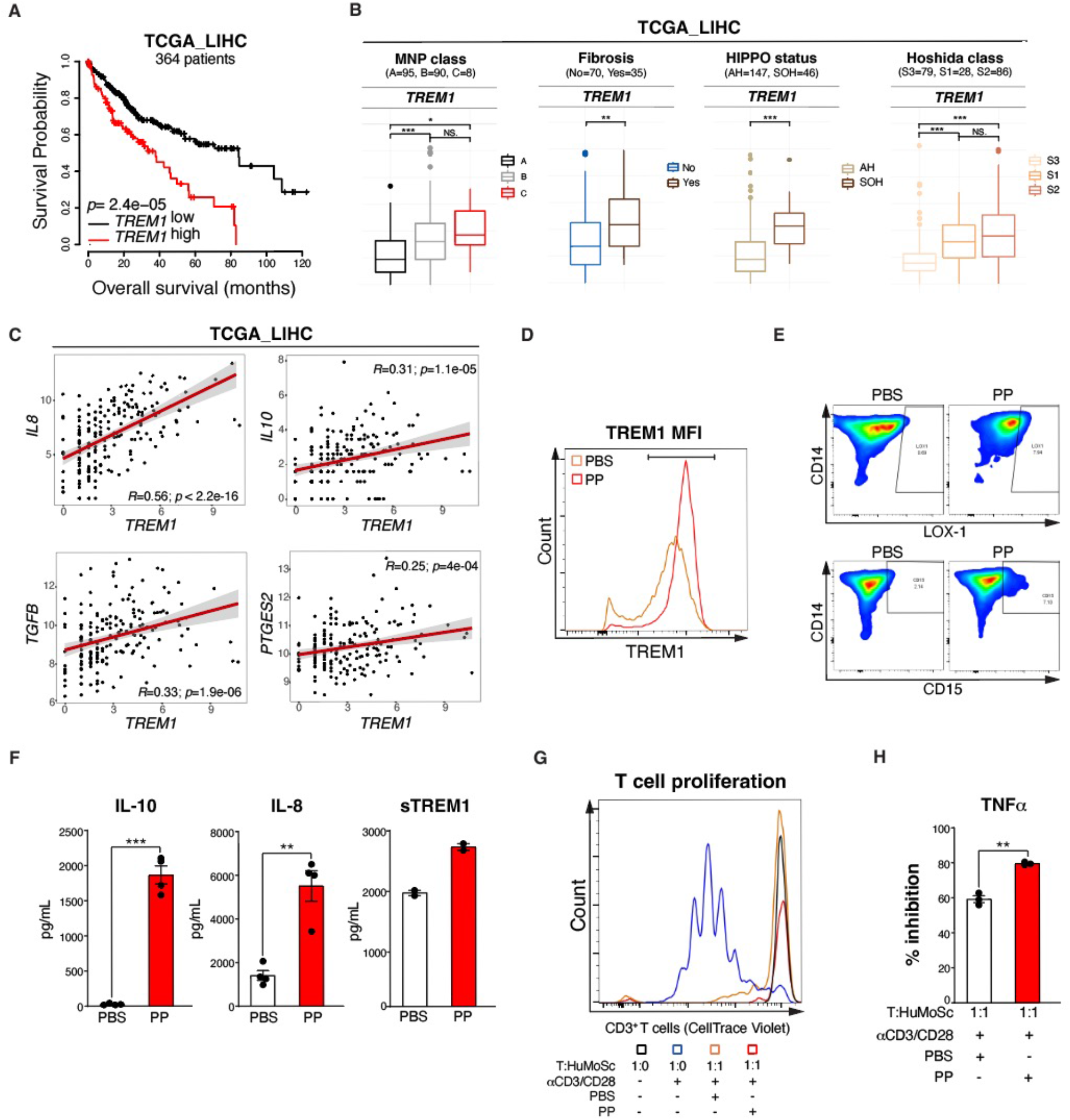
TREM1 associates with poor prognosis in solid tumors and promotes the immunosuppressive function of M_reg_ cells. (A) Kaplan-Meier survival curves of patients with HCC (n=364) (Menyhart *et al*., 2018) according to *TREM1* expression. (B) Box plots depicting the average expression of *TREM1* according to MNP infiltrate profiles, fibrosis, HIPPO status or Hoshida class. (C) Scatter plots depicting Pearson correlations between the expression of *TREM1* and genes associated with immunosuppression (*TGFB, IL10, IL8* and *PTGES2*). (D) Histograms depicting TREM1 expression on HuMoSc treated on day 6 of differentiation with PBS or a PP complex of peptidoglycan (PGN) and PGN recognition protein 1 (PGLYRP1) for 24 hours. (E) Representative FACS scatterplot depicting cells positive for the expression of CD14 (y- axis), LOX-1 (x-axis, top) and CD15 (x-axis, bottom) as in (D). (F) Bar graphs depicting the concentration of IL-10, IL-8 and soluble TREM1 (sTREM1) released from HuMoSc treated with PBS or PP as in (D). (G) Representative FACS histogram depicting CellTrace Violet dilution as a readout of CD3^+^ T cell proliferation for 6 days in the listed conditions (H) Quantification of the % inhibition of TNFα production by a CD3^+^ T cells stimulated or not with anti-CD3/CD28 in co-culture with HuMoSc pre-treated withf PBS or PP as in (D).

## Discussion

Here, investigation of the innate immunity hepatic cellular landscape of patients with HCC revealed previously unreported transcriptional and phenotypic states of different innate lymphoid and myeloid cells. Our work reports four novel findings with important clinical implications: 1) HCC-associated genesis of cells exhibiting mixed lymphoid and myeloid identities as well as cells with inter-myeloid mixed states, potentially stemming from emergency myelopoiesis and trained immunity responses to tumor growth (Mulder et al., 2019; Pietras et al., 2016; Sica et al., 2019); 2) a heterogeneity of myeloid states, yet with only specific subsets infiltrating the tumor; 3) identification of a TREM1^+^CD163^+^ M_reg_ subset with strong immunosuppressive activity that expands in the steatohepatitis setting, populates fibrotic lesions in HCC, and associates with poor prognosis in HCC; and 4) demonstration of a role of TREM1 in amplifying the immunosuppressive function of human monocyte-derived suppressor cells and promoting HCC growth *in vivo*.

The first observation applies particularly to a population of cytotoxic cells scoring highly for an NK signature yet segregating with myeloid cells in a UMAP visualization. Cells with similar properties have been previously described primarily in mouse models of cancer, and while they were initially thought to be DCs acquiring a ‘killer’ capacity (Chan et al., 2006; Taieb et al., 2006), they were later shown to derive from lymphoid progenitors in the bone marrow (Blasius et al., 2007; Caminschi et al., 2007; Vosshenrich et al., 2007) and renamed pre-mature NK (pre-mNK) (Guimont-Desrochers et al., 2012). Our results argue that myeNK are not pre-mNK for several reasons: a) trajectory analysis positioned MyeNK nearest to NK4 cells, which is a fully differentiated NK state; b) potency differentiation analysis similarly highlighted that both MyeNK and NK4 are the least entropic compared to other NK cells, particularly cycling NKs that in contrast has high entropy rates suggestive of a precursor state; and c) MyeNK were highly cytotoxic as demonstrated functionally in a cancer-killing assay, which is at odds with a pre-mature state. Our single cell analysis unravels the expansion of myeNK in the NASH setting, particularly in the juxta-tumoral tissue. Metabolic signals or inflammatory mediators induced by metabolic dysregulation might thus drive their genesis, recruitment to the liver and/or acquisition of myeloid features. This is in line with Theurich *et al*. who reported obesity-associated IL-6-dependent induction of an NK population with myeloid properties (Theurich *et al*., 2017). Cells expressing both cytotoxic and myeloid genes have also been reported in human blood (Villani et al., 2017) and HCC (Sun *et al*., 2021). However, in these studies, they were referred to as monocytes, without further characterization. We posit that MyeNK cells stem from an inflammatory signal amplified in the steatohepatitis setting. Whether the strong cytotoxic capacity of MyeNK is beneficial in NASH-HCC remains to be clarified.

Another phenomenon elicited in the tumor setting presumably through emergency myelopoiesis is the genesis and recruitment of MDSCs (Loftus et al., 2018). Our analysis demonstrates a significant heterogeneity in myeloid populations enriched in MDSC signature genes (Condamine *et al*., 2016), warranting reclassification of MDSCs according to more precise phenotypic and functional properties (Hegde *et al*., 2021). We show that *THBS1* expression marks myeloid cells enriched in MDSC genes (Condamine *et al*., 2016). They were found in three distinct states in HCC, including one with dual granulocyte- and macrophage/DC-lineage features, identified as *TREM1^+^CD163^+^*. Our results show that this subset exhibits a more potent immunosuppressive activity compared to other *THBS1_*M cells, strongly inhibiting T cell proliferation and effector functions *ex vivo*. We show, using spatial transcriptomics, that this subset highly populates fibrotic lesions in HCC in close associations with pro-fibrogenic and pro-tumorigenic fibroblast subsets. This subset was exclusively found in patients with steatohepatitis HCC in our cohort. Consistently, using *in vivo* models of liver disease, we observed a NASH-associated expansion of TREM1^+^ myeloid cells among the CD11b^+^LY6C^+^LYC6G^+^ population, which is consistent with the reported reliance of MDSC on cellular lipid metabolism (Veglia *et al*., 2021) and upregulated cell surface expression of lipid receptors and transporters, notably the oxidized low-density lipoprotein (oxLDL) receptors CD36 and LOX-1 (*OLR1*) and the fatty acid transporter FATP2 (Veglia et al., 2019). This was accompanied by notable TREM1 induction on this subset in NASH, that was not observed in control mice and to a much lower extent in liver injury induced by CCL4 treatment. In line with a recent report demonstrating progressive accumulation of CD8^+^TNF^+^PD-1^+^ T cells in the NASH-affected liver that exacerbates HCC in a mouse model in a CD8^+^ T cell- and TNF- dependent manner (Pfister *et al*., 2021), we observed enhanced hepatic infiltration of CD8^+^TNF^+^ T cells in NASH, but this was not unique to this condition, as it was also observed in CCL4-treated mice. Thus, besides CD8^+^TNF^+^ T cell infiltration, NASH also elicits TREM1^+^ M_reg_ recruitment, which might underlie resistance to immunotherapy.

TREM1, a cell surface receptor expressed on monocyte-derived inflammatory macrophages and neutrophils, has been previously described as an amplifier of inflammation (Bouchon et al., 2001). It was initially proposed to promote HCC development through pro-inflammatory cytokine production by Kupffer cells (Wu et al., 2012). However, this early analysis used a non-discriminatory staining that marks a heterogeous group of myeloid cells, preventing precise identification of deleterious myeloid cell subsets. Our scRNA-seq data reveal that TREM1 is a discriminatory feature of *THBS1_*M cells distinguishing them from other MNPs including monocytes, macrophages/TAMs and DCs in the liver of HCC patients. Among these, we identify a *TREM1^+^CD163^+^*M_reg_ subset as a potent immunosuppressive population that associates with poor prognosis in HCC. We show that *TREM1* expression *per se* is predictive of poor clinical outcomes in several solid tumors and with ICB resistance. TREM1 has been implicated in mediating liver fibrosis (Nguyen-Lefebvre et al., 2018) and immunotherapy resistance in a mouse model of liver cancer (Wu *et al*., 2019). However, the underlying mechanisms have not been fully explored. Functionally, we demonstrate that TREM1 ligation by cognate ligand on human monocyte-derived suppressor cells promotes their acquisition of features linked to a suppressive phenotype, such as enhanced expression of LOX- 1, CD15 and IL-10, and boosts their capacity to suppress T cell proliferation and effector functions.

Collectively, our study supports the stratification of patients according to HCC etiology to define optimal therapeutic regimens, and points to TREM1 targeting as an attractive therapeutic option in NASH-associated HCC.

## Authors’ contributions

All authors revised the manuscript and approved the final version.

## Acknowledgements

We thank the TBMCore flow cytometry platform operator Vincent Pitard, Marlène Maître at the Magendie Neuro center, Sébastien Marais at the Bordeaux Imaging Center, Aurélia Le Dantec, Gaël Galli and Sébastian Lillo for help and advice. We also thank the Bordeaux Bioinformatic Center (CBiB) for computational resources. This work is funded by operating grants to MS from the ARC foundation, IDEX Bordeaux and ITMO cancer and an infrastructure grant to MS from the New Aquitaine region. JG and DC are funded by the ARC foundation grant and ER received a studentship from the SIRIC BRIO. JG, DC and ER also received salary funding from the New Aquitaine region.

## SUPPLEMENTAL FIGURES AND TABLES

**Figure S1. Related to Figure 1.**
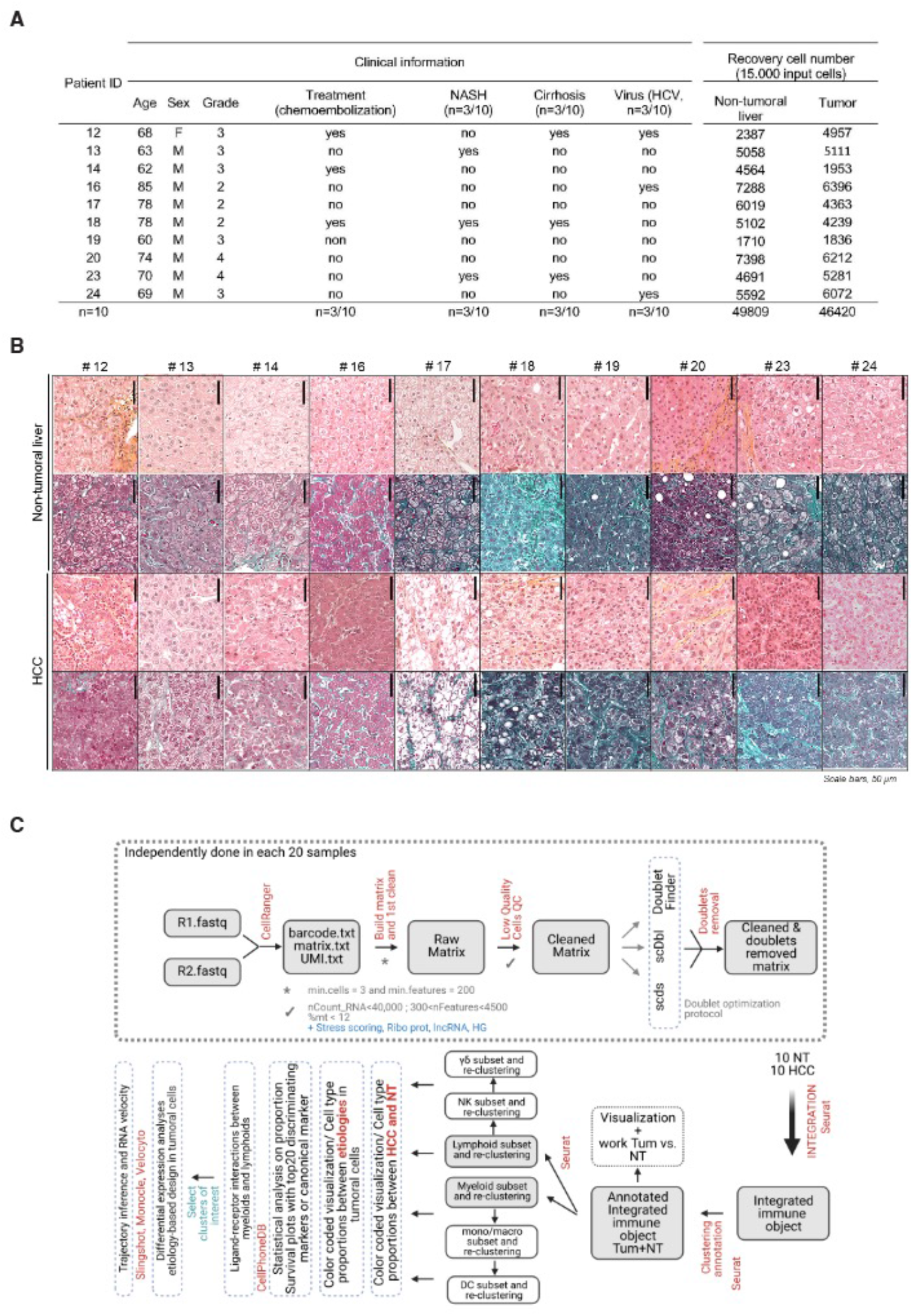
(A) Clinical characteristics of the patients included in this study. Complete information is provided in Table S1. (B) HES staining of NT and tumoral tissue sections from all patients in the study. (C) Pipeline of scRNA-seq bioinformatic analyses

**Figure S2.**
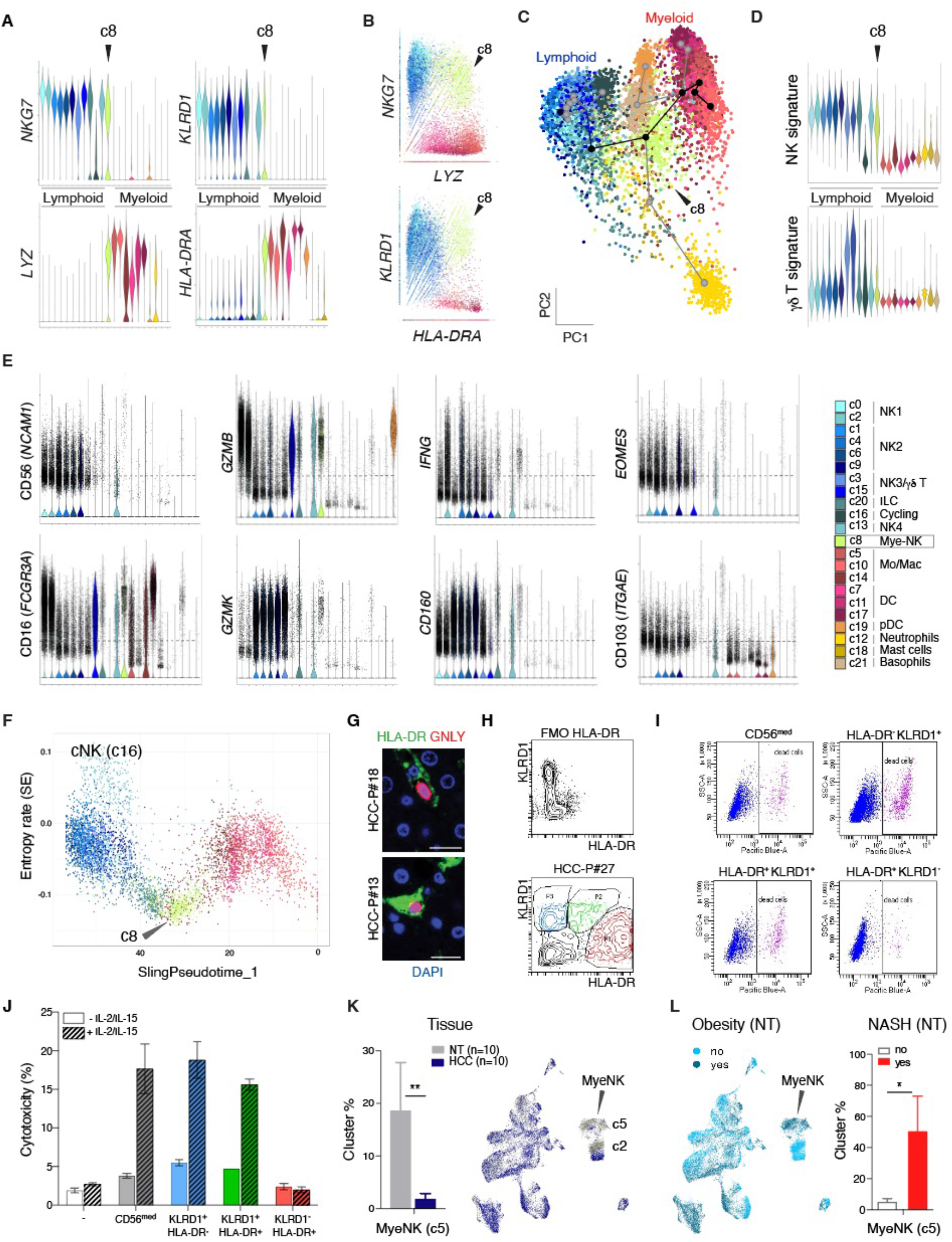
Identification of a MyeNK population in HCC depleted from the tumor and associated with good prognosis. (A) Violin plots depicting the expression levels of lymphoid (*NKG7, KLRD1*) and myeloid (*LYZ, HLA-DRA*) features in the different innate immunity main clusters. (B) Scatter graphs depicting clusters containing single versus double positive cells for the expression of *NKG7* and *LYZ* or *KLRD1* and *HLA-DRA*. (C) Slingshot trajectory analysis identifying MyeNK at the intersection between myeloid and lymphoid lineages. (D) Violin plots depicting the scores of NK versus γδ T cell signatures in the different innate immunity main clusters. (E) Violin plots depicting the expression levels of NK features in the different innate immunity main clusters. (F) Scatterplot of signaling entropy (SE) against Slingshot Pseudotime values. Low SE is indicative of a differentiated state. (G) Multiplex immunofluorescence staining of HLA-DR (green) and GNLY (red) expression; DAPI marks nuclei (blue). (H) Representative FACS scatterplot depicting cells positive for the expression of HLA-DR (red), KLRD1 (blue) and HLA-DR^+^ KLRD1^+^ double positive cells (green). FMO, fluorescence minus one. (I) Scatterplots depicting Pacific Blue-positive dead K562 cells killed by NK2 (CD56^med^), NK1 (KLRD1^+^ HLA-DR^-^) or MyeNK (KLRD1^+^ HLA-DR^+^) cells isolated by FACS sorting from liver of human patients with HCC (n=3), in *in vitro* co-culture experiments. (J) Quantification of dead cells from the *in vitro* killing assay, as in (H). One of two representative experiments is shown. Data represent the mean +/- SEM. (K) Left, MyeNK (c5) frequencies in the adjacent NT versus HCC of the 10 patients investigated; right, UMAP colored by tissue site, HCC (blue) and NT (grey). **p<0.01 using Mann-Whitney tests. (L) Left, UMAP of cells in NT colored by obesity status, obese (dark blue), non-obese (light blue); right, MyeNK (c5) frequencies in NT according to NASH status. *p<0.05 using Mann-Whitney tests.

**Figure S3. Related to Figure 2.**
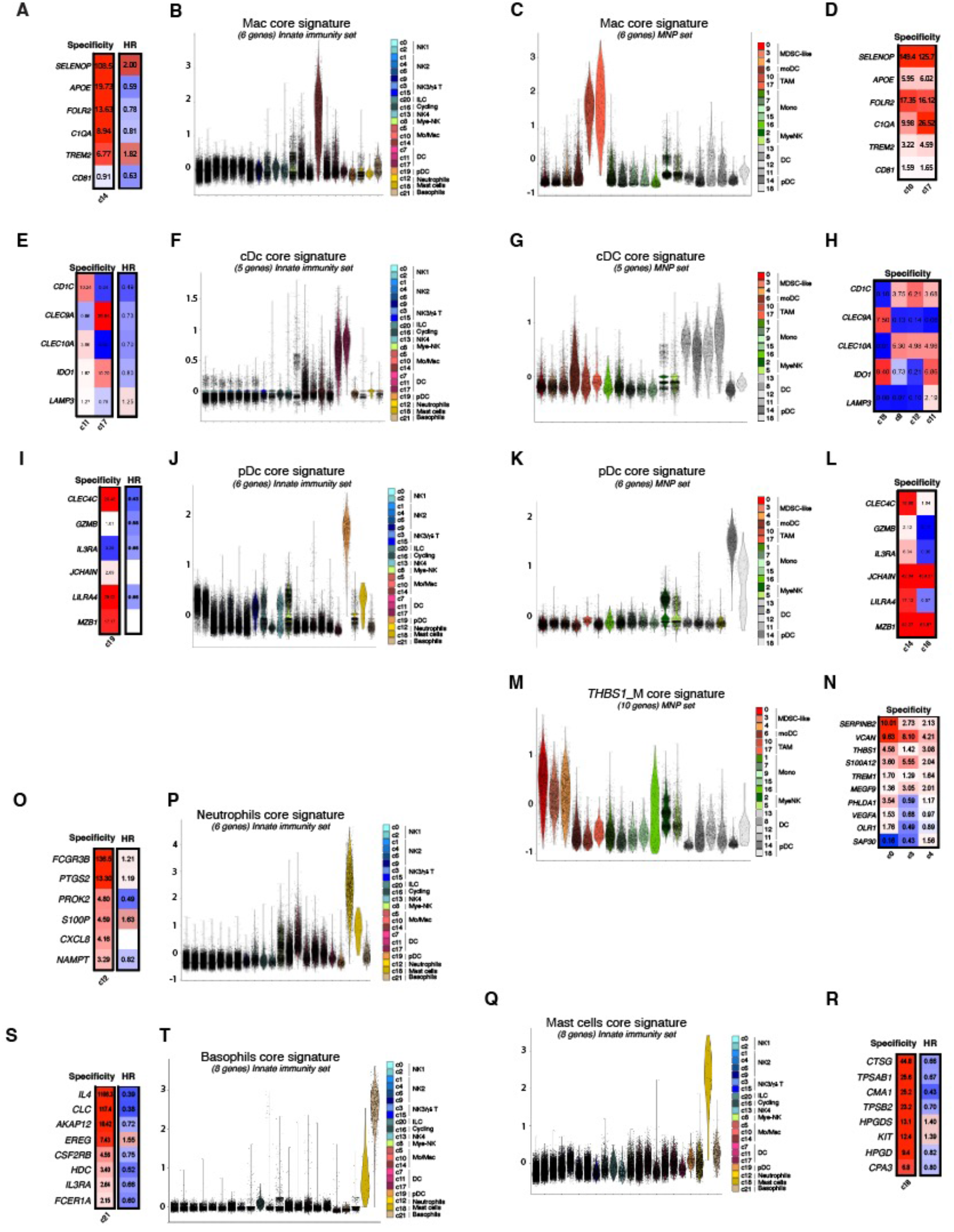
(A) Core Mac signature specificity and association with mOS in patients with HCC. HR, hazard ratio. (B) Violin plots depicting the score of Mac core signature in the different innate immunity main clusters. (C) Violin plots depicting the score of Mac core signature in the different MNP clusters. (D) Core Mac signature specificity applied to the two Mac clusters of the MNP dataset (c10, 17) and association with mOS in patients with HCC. HR, hazard ratio (E) Core cDC signature specificity and association with mOS in patients with HCC. HR, hazard ratio. (F) Violin plots depicting the score of cDC core signature in the different innate immunity main clusters. (G) Violin plots depicting the score of cDC core signature in the different MNP clusters. (H) Core cDC signature specificity applied to the four cDC clusters of the MNP dataset (c8, 11, 12, 13) and association with mOS in patients with HCC. HR, hazard ratio (I) Core pDC signature specificity and association with mOS in patients with HCC. HR, hazard ratio. (J) Violin plots depicting the score of pDC core signature in the different innate immunity main clusters. (K) Violin plots depicting the score of pDC core signature in the different MNP clusters. (L) Core pDC signature specificity applied to the two pDC clusters of the MNP dataset (c14, 16) and association with mOS in patients with HCC. HR, hazard ratio (M) Violin plots depicting the score of *THBS1_*M core signature in the different MNP clusters. (N) Core *THBS1_*M signature specificity applied to the three *THBS1_*M clusters of the MNP dataset (c0, 3, 4) and association with mOS in patients with HCC. HR, hazard ratio (O) Core neutrophils signature specificity and association with mOS in patients with HCC. HR, hazard ratio. (P) Violin plots depicting the score of neutrophils core signature in the different innate immunity main clusters. (Q) Core mast cells signature specificity and association with mOS in patients with HCC. HR, hazard ratio. (R) Violin plots depicting the score of mast cells core signature in the different innate immunity main clusters. (S) Core basophils signature specificity and association with mOS in patients with HCC. HR, hazard ratio. (T) Violin plots depicting the score of basophils core signature in the different innate immunity main clusters.

**Figure S4. Related to Figure 2.**
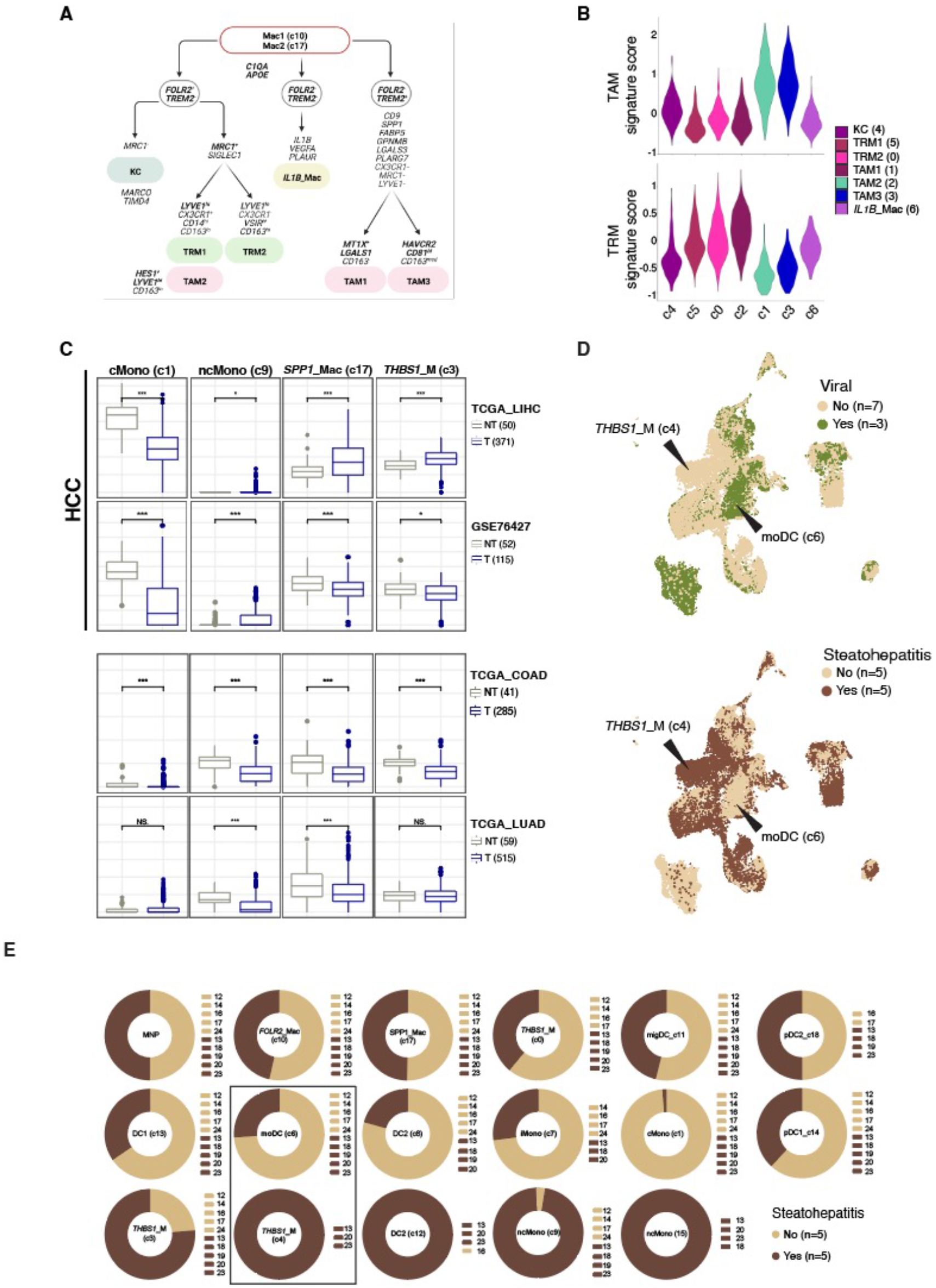
(A) A schematic illustration summarizing marker gene expression in the different macrophage subsets in HCC; TAMs (pink), TRMs (green). (B) Violin plots depicting the score of TAM versus TRM signatures (C) Box plots depicting the predicted proportions of MNP subsets by CIBERSORTx deconvolution of bulk transcriptomics datasets from four cohorts of patients, with either HCC (TCGA_LIHC; GSE76427), colon adenocarcinoma (TCGS_COAD) or lung adenocarcinoma (TCGA_LUAD). (D) UMAP of cells in HCC+NT colored by viral (green) or steatohepatitis (brown) positive status. (E) Donut plot depicting MNP cell proportions in HCC according to steatohepatitis positive status (brown).

**Figure S5. Related to Figure 3.**
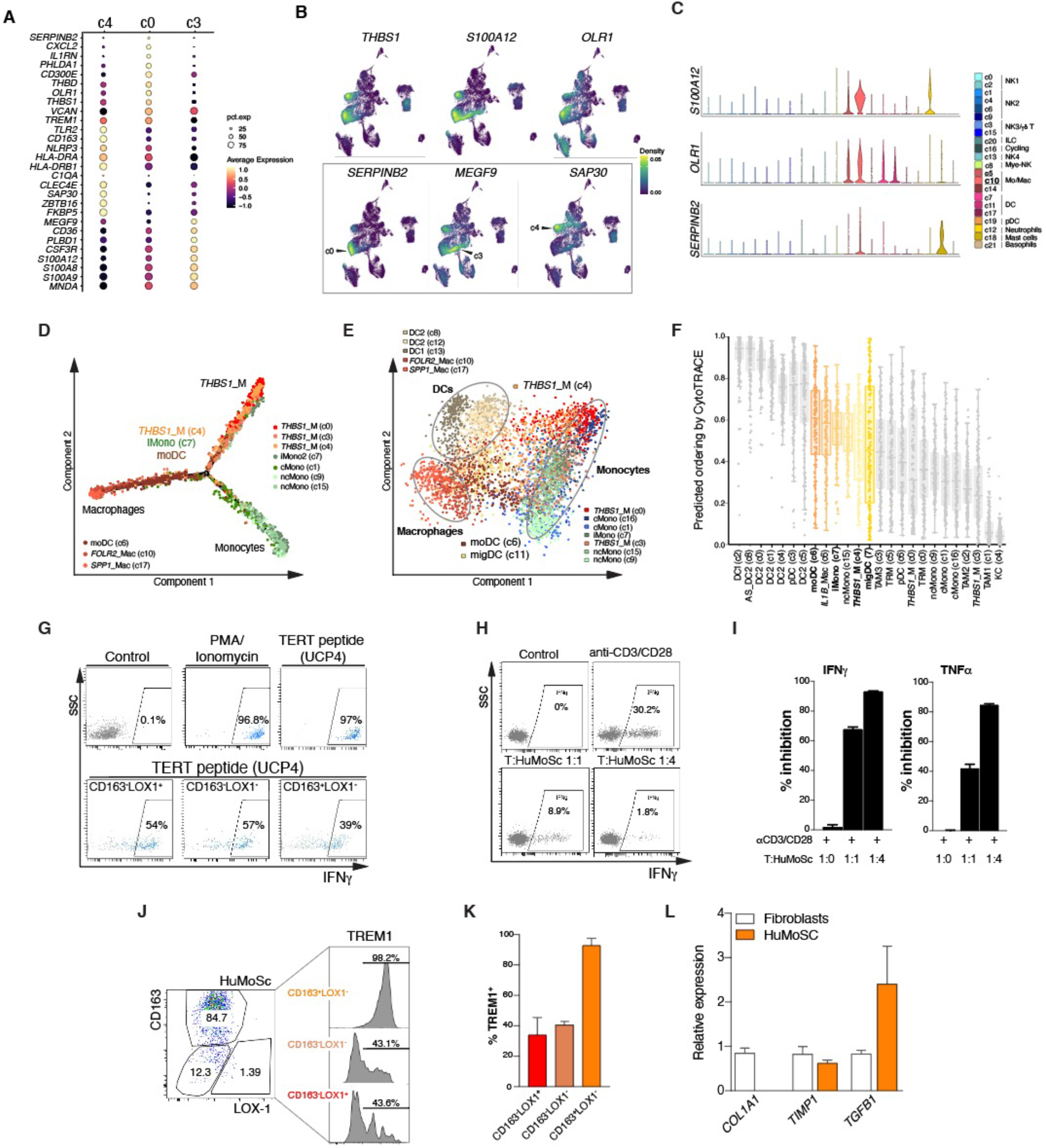
(A) Dot plot depicting average expression per cluster of top discriminating genes. (B) Density plots representing the expression of discriminating genes marking the three *THBS1_*M clusters universally (top) or selectively (bottom). (C) Violin plots depicting the expression levels of discriminating genes in the different innate immunity main clusters. (D) Pseudotime trajectory analysis using Monocle2 of the three *THBS1_*M clusters together with those encompassing macrophages (c10, 17), moDC (c6) and monocytes (c1, 7, 9, 15, 16) from the MNP dataset. (E) Principal Component Analysis of 300 cells from all MNP clusters. (F) Predicted ordering of differentiation by CytoTRACE analysis (G) Representative FACS scatter plots depicting intracellular IFNγ staining in TERT-specific CD4^+^ T cells stimulated or not with PMA/ionomycin or a TERT peptide (UCP4) in the presence or absence of myeloid cells with differential expression of TREM1 and LOX-1 (H) Representative FACS scatter plots depicting intracellular IFNγ staining in CD3^+^ T cells stimulated or not with anti-CD3/CD28 in the presence or absence of HuMoSc at the indicated co-culture ratios. (I) Quantification of the % inhibition of IFNγ and TNFα production by CD3^+^ T cells stimulated or not with anti-CD3/CD28 in the presence or absence of HuMoSc at the indicated co-culture ratios. (J) Phenotypic characterization of HuMoSc using the gating strategy in (Figure 3L), and histograms depicting TREM1 expression among the different subsets. (K) Quantification of the frequencies of TREM1^+^ cells among the different HuMoSc gates shown in (J). (L) Expression of *COL1A1, TIMP1* and *TGFB1* in HuMoSc and fibroblasts cultures, measured by quantitative RT-PCR. Note that *COL1A1* is not expressed in HuMoSc.

**Figure S6, related to Figure 5.**
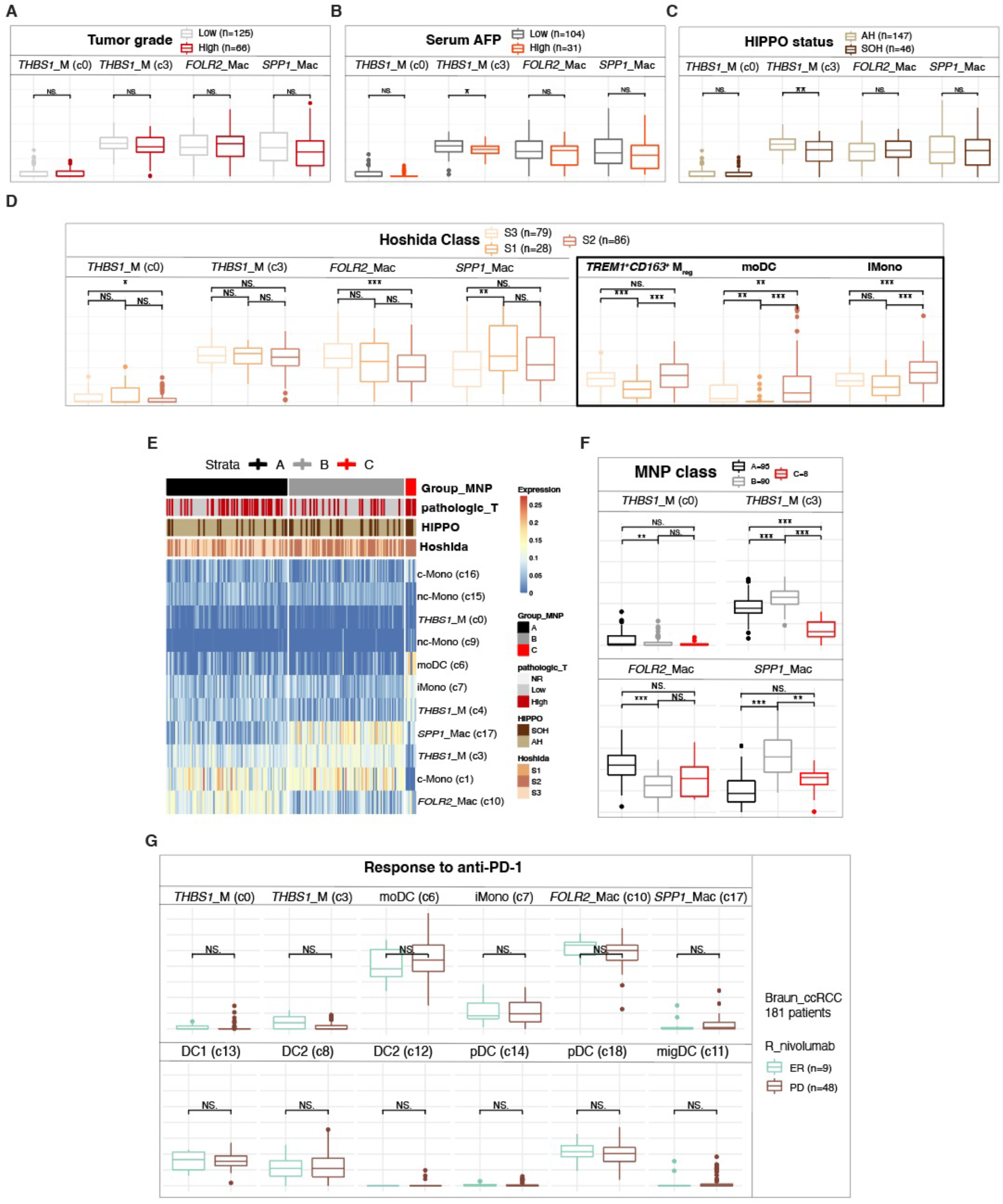
(A-D) Box plots depicting the predicted proportions of MNP subsets by CIBERSORTx deconvolution of bulk transcriptomics data from the TCGA_LIHC cohort according to tumor grade (A), serum AFP (B), HIPPO status (C) or Hoshida class (D). (E) Heatmap depicting estimated proportions of MNP subsets derived from CIBERSORTx in HCC patients from the TCGA_LIHC cohort (F) Box plots depicting the predicted proportions of MNP subsets according to patient group from (E). (G) Box plots depicting the predicted proportions of MNP subsets according to patient response to anti-PD-1 (Nivolumab) in a cohort of patients with clear cell renal cell carcinoma (ccRCC).

**Figure S7, related to Figure 6.**
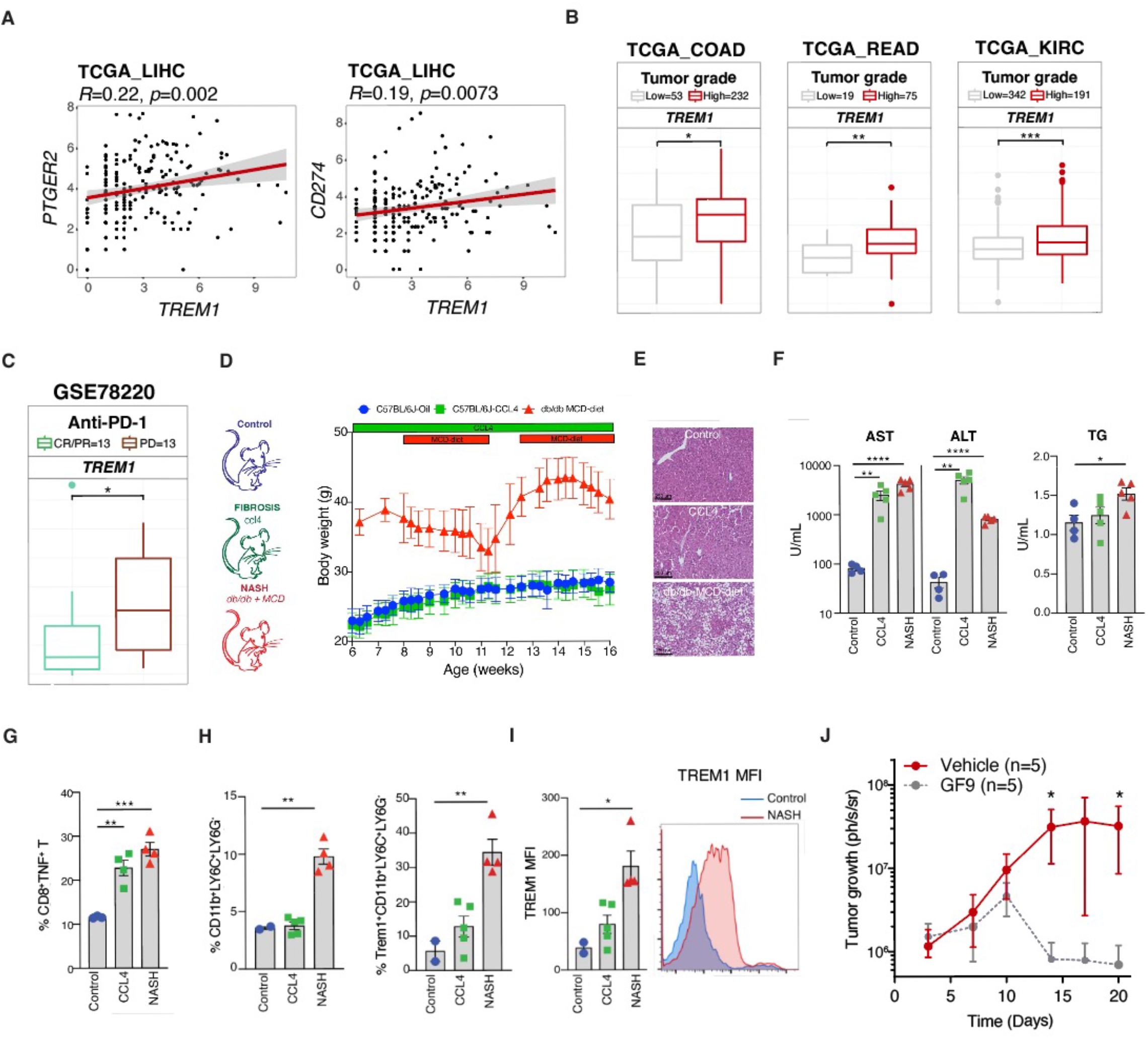
(A) Scatter plots depicting Pearson correlations between the expression of *TREM1* and *PTGER2* (left) or *CD274* (right). (B) Box plots depicting *TREM1* expression according to tumor grade in three cohorts of solid tumors, TCGA_COAD (colon adenocarcinoma), TCGA_READ (rectum adenocarcinoma) and TCGA_KIRC (kidney renal cell carcinoma). (C) Box plots depicting *TREM1* expression according to patient response to anti-PD-1 in a cohort of patients with metastatic melanoma. (D) Study design for the mouse models of NASH and fibrosis and mouse body weight from a representative experiment. (E) H&E staining of liver tissue sections from mice of the indicated condition. Scale bar, 250μm. (F) AST, ALT and triglycerides (TG) serum concentration in C57BL/6J mice treated with CCL4 (green) and db/db mice fed a MCD-diet (red). Age-matched C57BL6J mice were used as controls (blue) (G) Percentage of CD8^+^TNF^+^ cells among CD3^+^ T cells. (H) Percentage (%) of CD11b^+^LY6C^+^LY6G^-^ (left panel) and TREM1^+^CD11b^+^LY6C^+^LY6G^-^ (middle panel). (I) Mean fluorescence intensity (MFI) quantification (left panel) and a representative histogram (right panel) of TREM1 in TREM1^+^CD11b^+^LY6C^+^LY6G^-^ in the indicated mouse groups. (J) Tumor volume by luminescence quantification in a liver orthotopic transplantable model of HCC using Hep55.1C cell-line transduced with a lentivirus encoding luciferase. Mice were treated with a vehicle control peptide or the GF9 peptide (i.p.) three times a week at 25 mg/kg starting on day 7 post tumor cells inoculation.

**TABLE S1.**
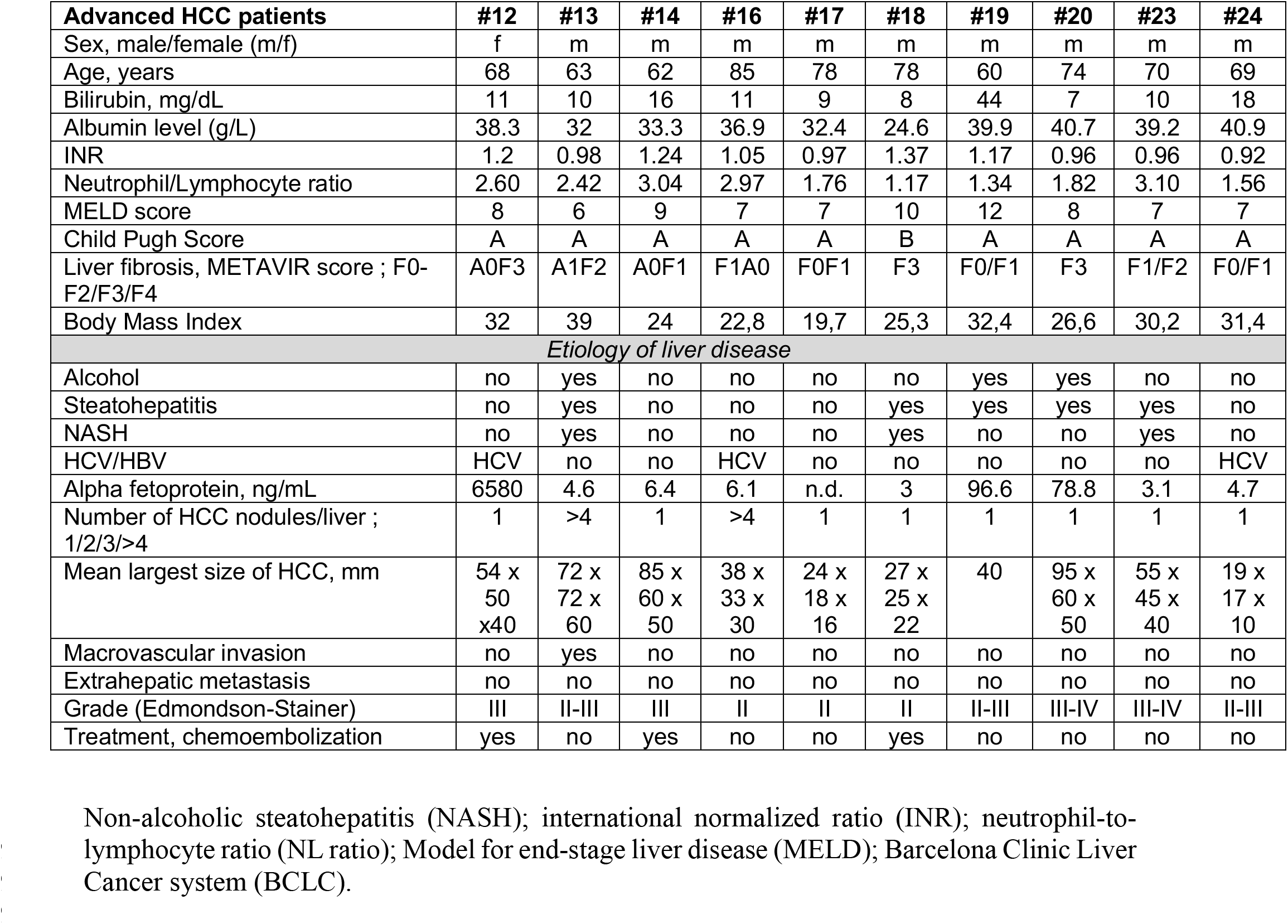
Patient Characteristics

## SUPPLEMENTAL TABLES (SEPARATE EXCEL FILES)

**TABLE S2. CELL NUMBER PER CLUSTER AND SUBCLUSTER**

**TABLE S3. PANGLAO LISTS**

**TABLE S4. DISCRIMINATING GENES LIST_TOP 100**

**TABLE S5. SIGNATURES**

**TABLE S6. THBS1_M DEGs**

**TABLE S7. VISIUM WEB SUMMARY**

**TABLE S8. DATA PROCESSING SUMMARY**

## STAR METHODS

### 1) KEY RESOURCES

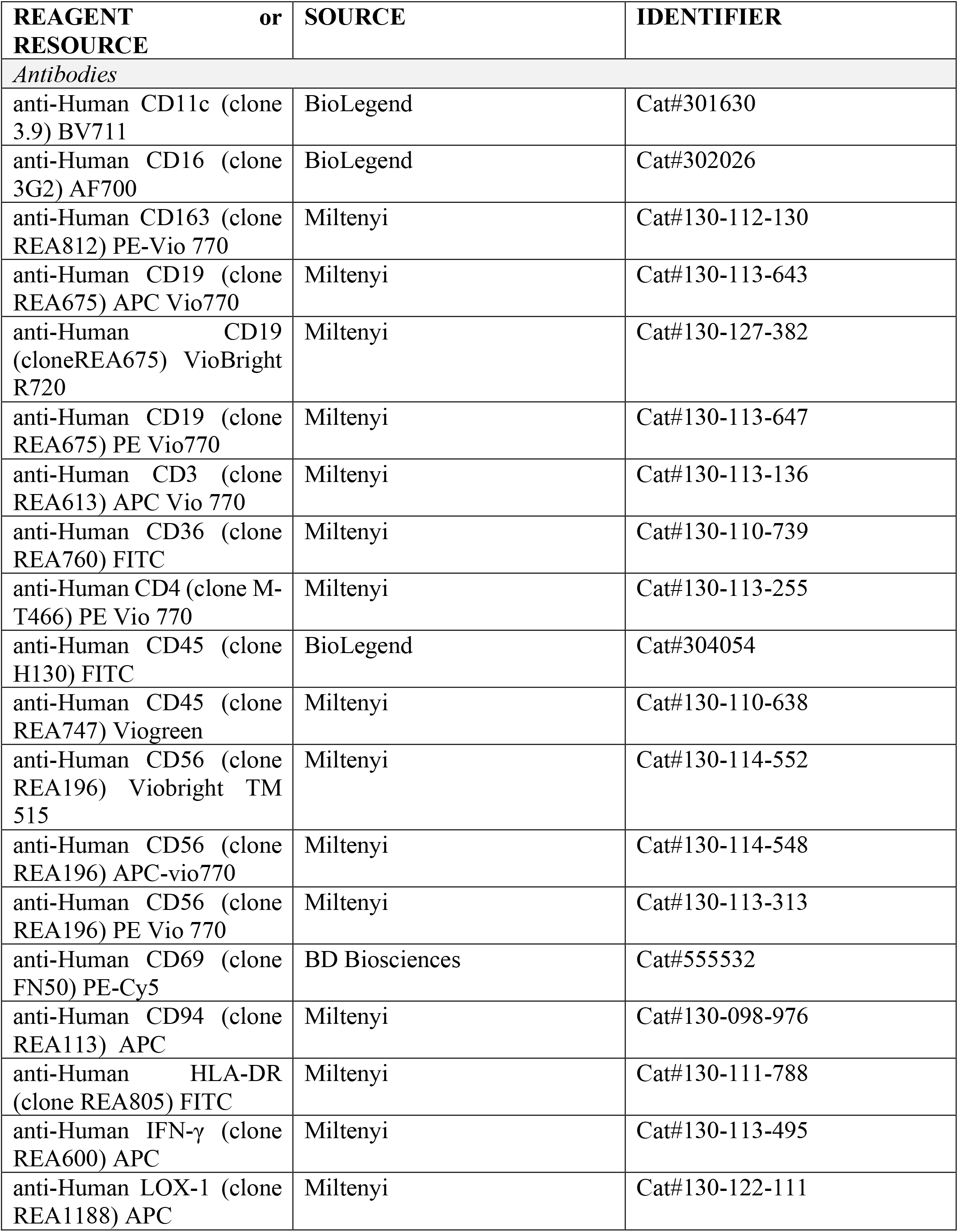
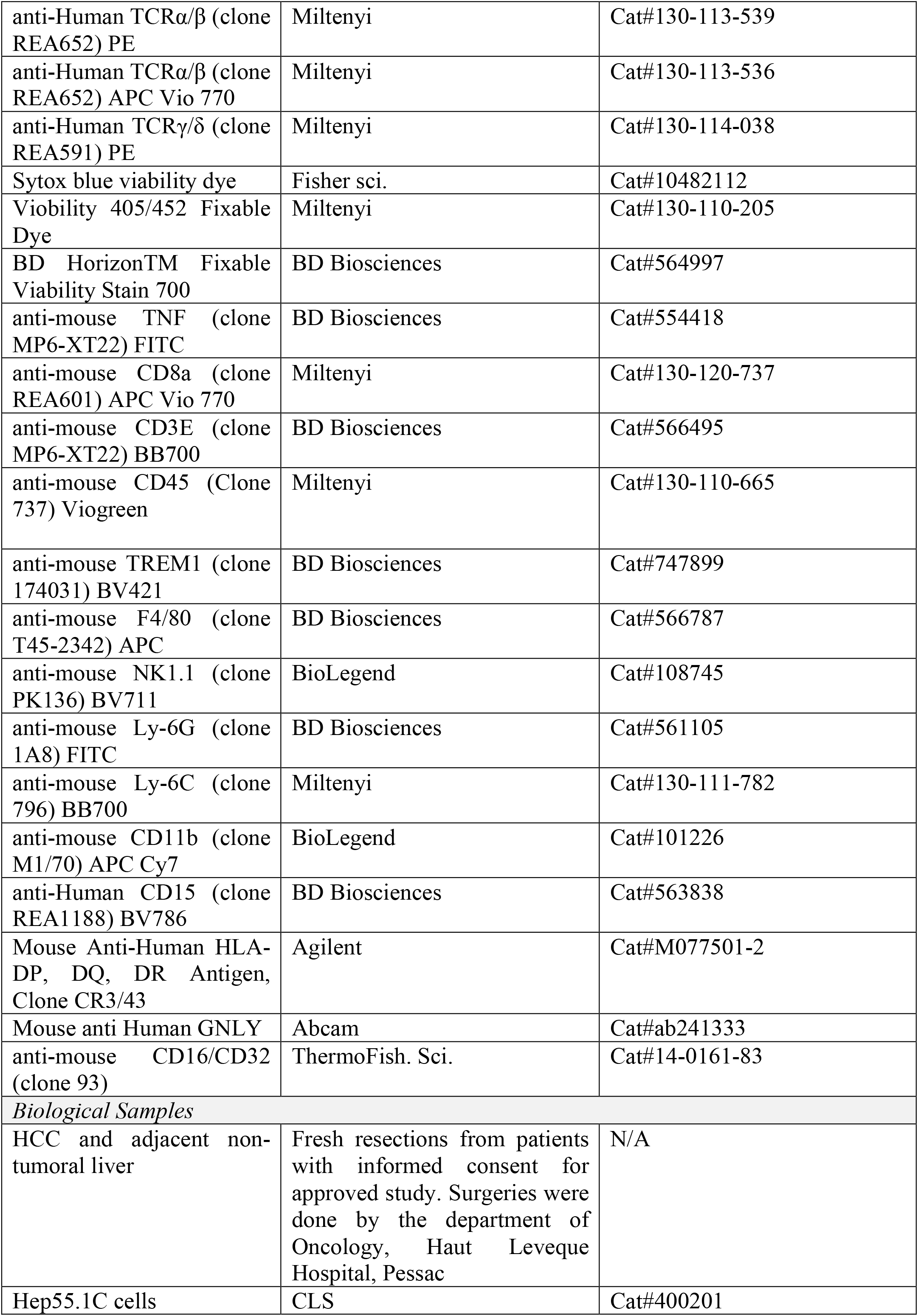
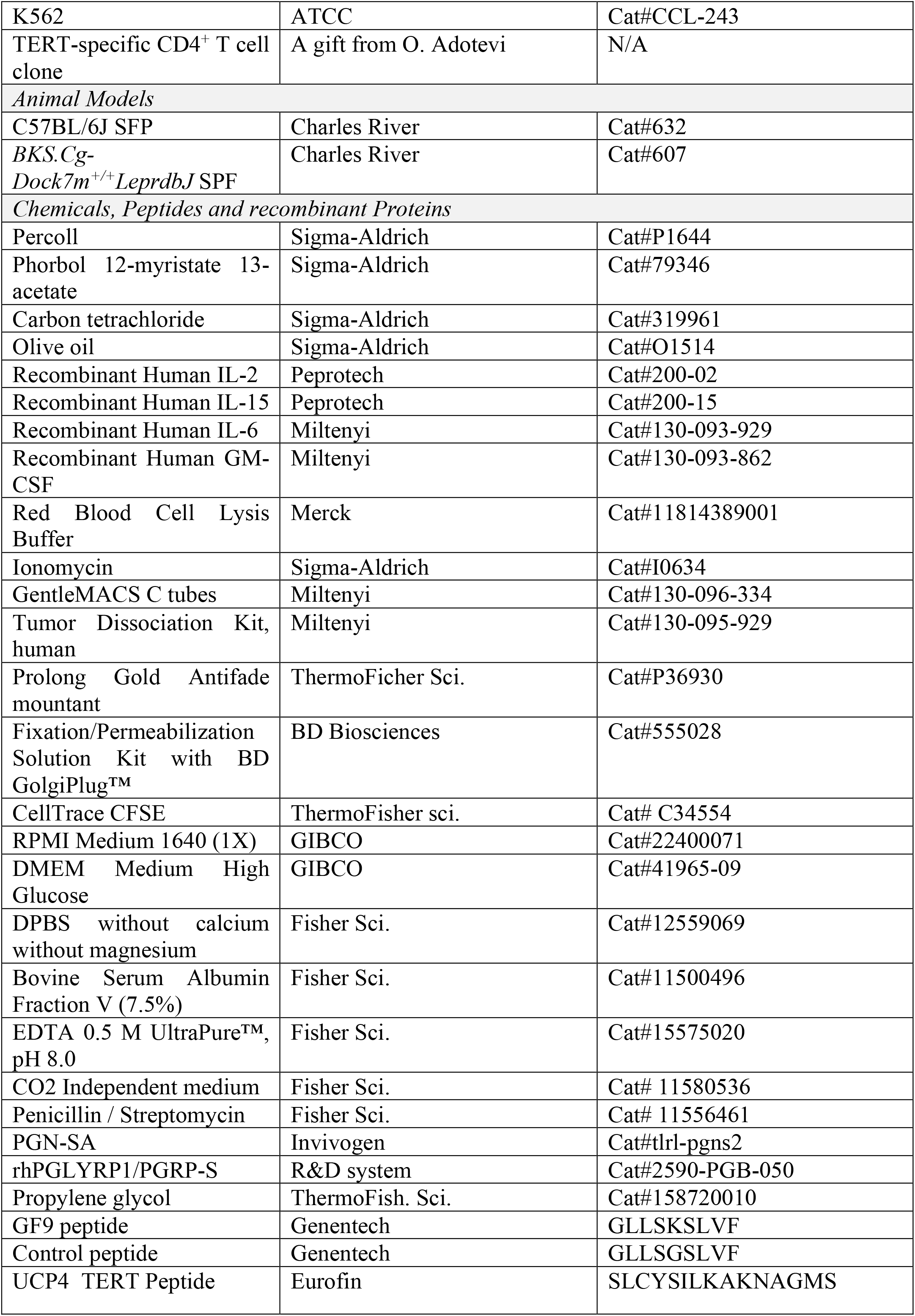
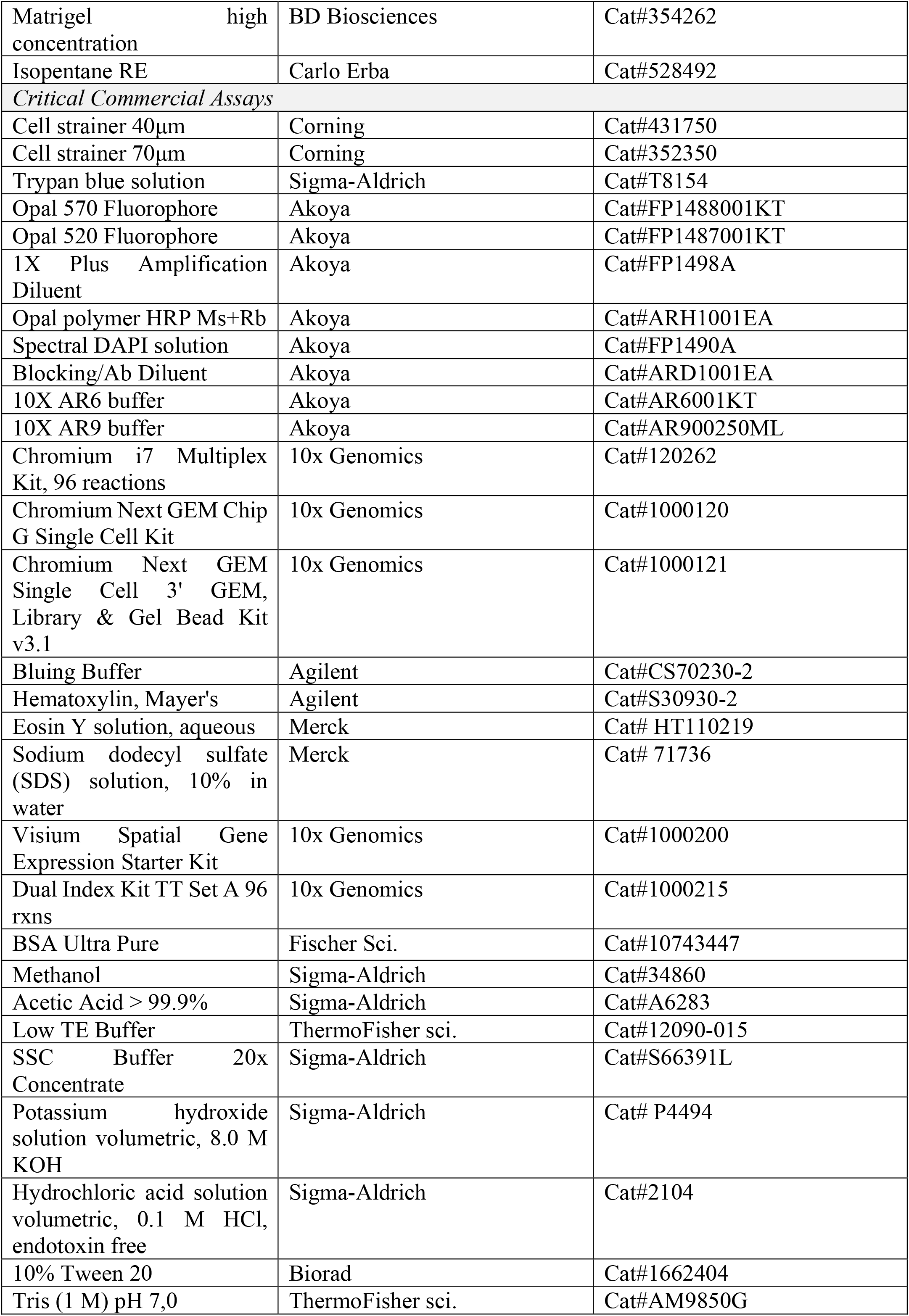
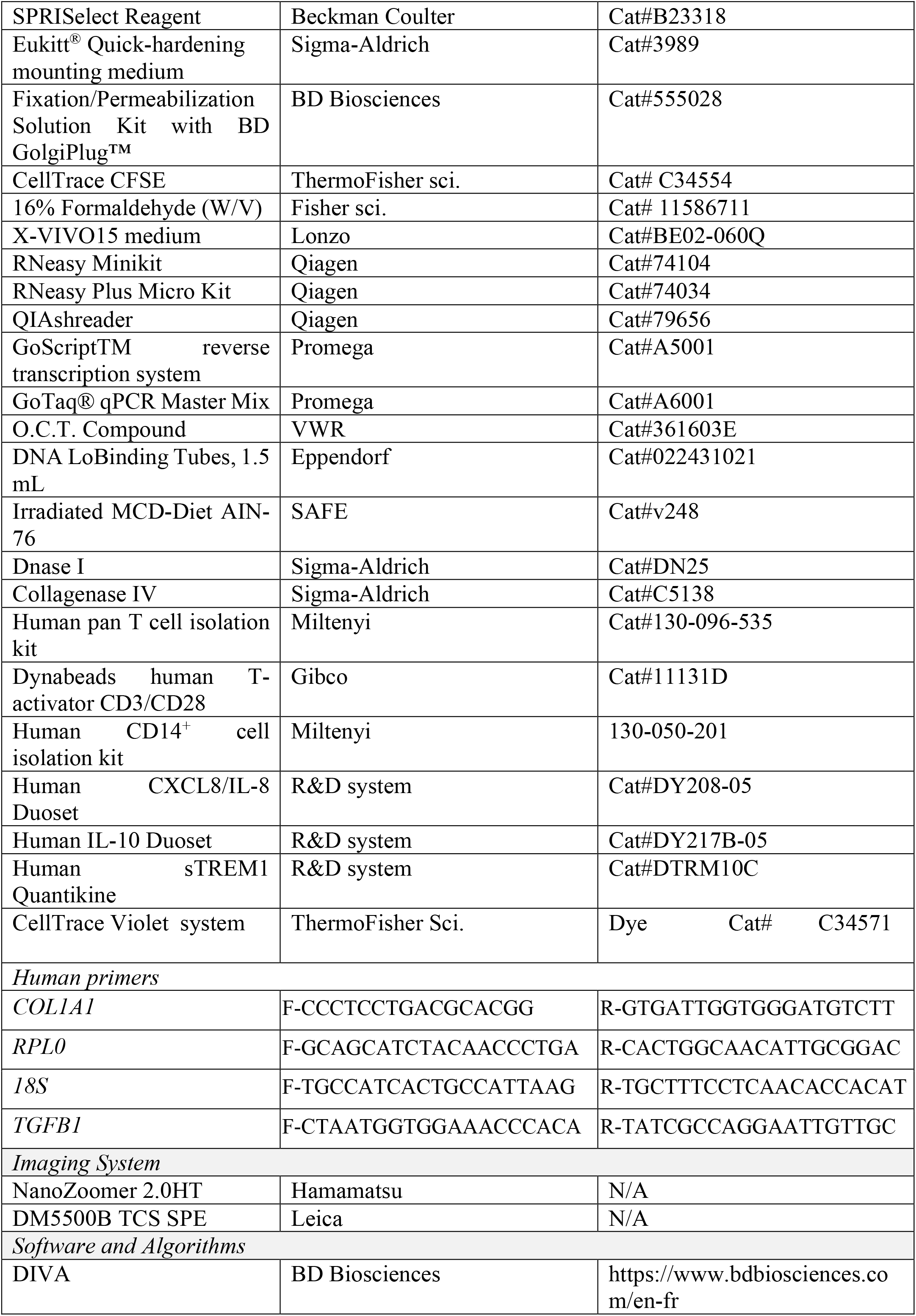
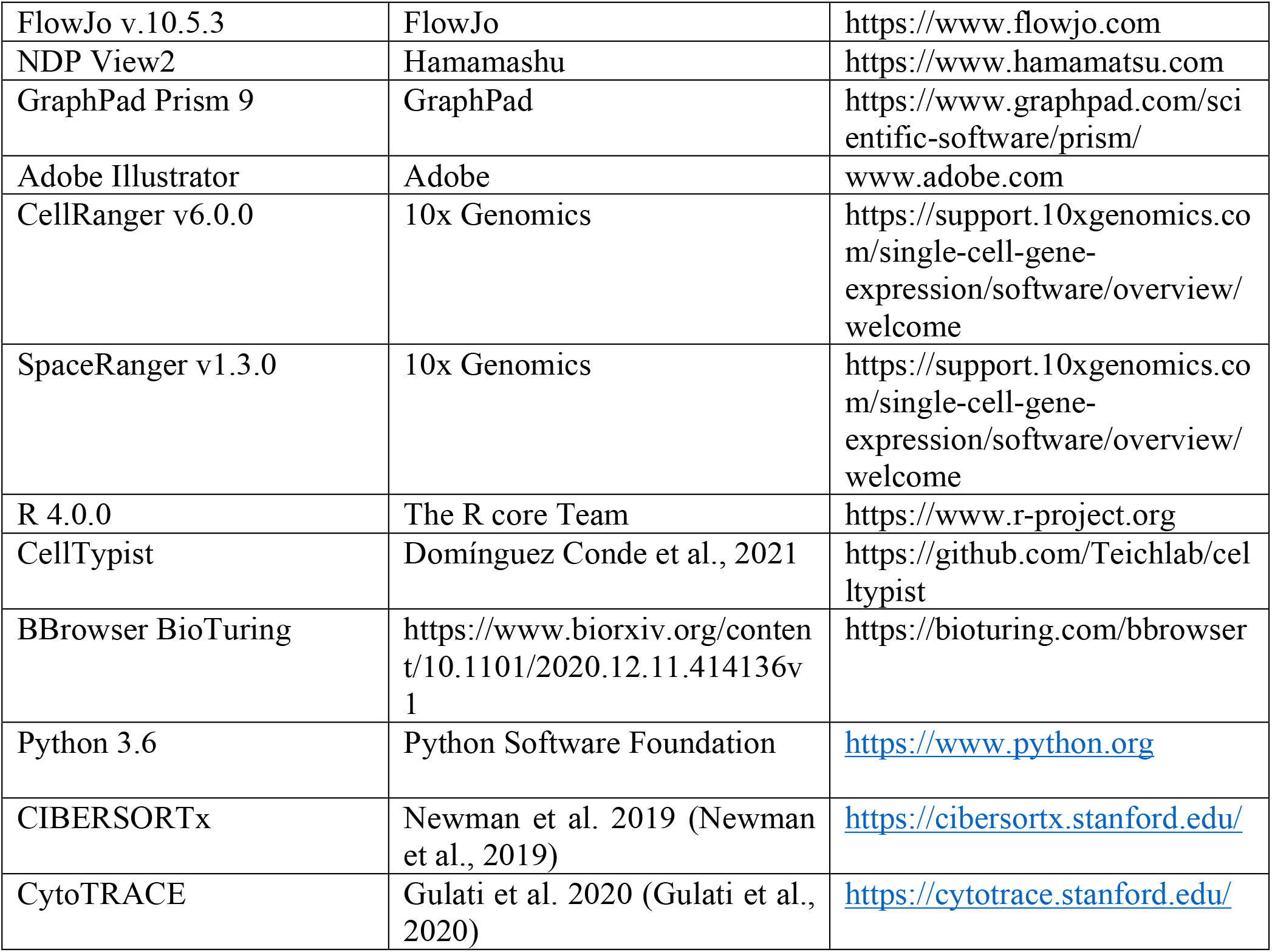

### 2) RESOURCE AVAILABILITY

#### Lead Contact

Further information and requests should be directed to and will be fulfilled by the Lead Contact, Dr. Maya Saleh.

### 3) EXPERIMENTAL MODELS AND SUBJECT DETAILS

#### Patient Clinical Characteristics

Samples were collected from patients diagnosed with HCC. Among the patients used for scRNAseq, five were obese with an IMC > 30, six were diagnosed as lower stage (stages <3) and four as higher stage (stages ≥3) according to Edmondson-Stainer histopathological classification. Three patients were HCV-infected, three had NASH, three were cirrhotic and three where alcoholic. The patients detailed clinical characteristics are summarized in Table S1.

#### Human HCC Samples

The adjacent non-tumoral (NT) and tumoral (HCC) tissues were obtained from patients undergoing liver resection surgery at the Haut Leveque Hospital (Pessac, France) with written consent.

#### Cell-Lines

The K562 human cell-line was obtained from the American Type Culture Collection and grown in RPMI 1640 supplemented with fetal bovine serum (FBS, 10%), glutamine (1%) and penicillin/streptomycin (1%) (named complete medium). Hep55.1C murine syngenic cell-line was obtained from the American Type Culture Collection and grown in DMEM, high glucose, GlutaMAX™ Supplement, pyruvate (Fisher 11594446) supplemented with fetal bovine serum (FBS, 10%) and penicillin/streptomycin (1%). These cell-lines were grown at 37°C in a humidified chamber with 5% CO2 and regularly tested as negative for mycoplasma. UCP4-specific CD4^+^ T cells were cultured as previously reported (Dosset et al., 2012).

#### Generation of HuMoSCs

HuMoSCs were generated according to a published protocol (Janikashvili *et al*., 2015). Briefly, healthy donor peripheral blood mononuclear cells (PBMC) were obtained from buffy coats by the means of Ficoll density gradient centrifugation. Monocytes were then isolated from PBMC by magnetic cell sorting (human CD14^+^ cell isolation kit; Miltenyi Biotec, 130-050-201). HuMoSCs were generated by incubating monocytes (1.10^6^ cells/mL) in RPMI 1640 supplemented with 10% FBS, 1% glutamine, pyruvate, Hepes, AANE and recombinant human GM-CSF (10 ng/mL; Militenyi Biotec, 130- 093-862) and IL-6 (10 ng/mL; Militenyi Biotec, 130-093-929) for 7 days. Sixty percent of the medium was replaced every 3 days. PGN/PGLYRP1 complex (2 µg/mL and 1 µg/mL, respectively) or PBS were added in the HuMoSC medium on day 6 for 24 hours. sTREM1, IL- 8 and IL-10 were quantified in the medium of HuMoSC cultures on day 7.

#### *In Vivo* Animal Studies

The mice used in this study were male C57BL/6J and BKS.Cg- *Dock7^m^+/+Lepr^dbJ^* purchased from Charles River. Mice were housed under specific pathogen-free conditions at the animal facility at Pessac (University of Bordeaux). All experimental procedures were approved by the local ethical committee in accordance with the regulations of the French ministry.

### 4) EXPERIMENTAL METHODS

#### Fibrosis and NASH mouse models

Liver fibrosis was induced in 6-week-old C57BL/6J male mice with intraperitoneal (i.p.) injection of CCL4 at the dose of 2µl/g body weight diluted 1:19 in olive oil, twice a week for 10 weeks. Adult 8-week-old male db/db mice were fed a Methionine- and Choline-Deficient diet (MCD-diet) from week 8 to week 16. The diet was halted 9 days between weeks 11.3 and 12.7. Comparison was made to age-matched olive-oil treated mice (controls, i.p. injection) and mice were sacrificed at 16-weeks of age.

#### Mouse liver dissociation

Mouse liver was dissociated in Gentle MACS C tubes containing RPMI medium supplemented by 1% glutamine, 1% P/S, 50 μl/mL DNAse I and 1 mg/mL collagenase IV for 45 minutes at 37°C under agitation. C tubes were run in GentleMACS dissociator before and after incubation (program mLiver_3, Miltenyi). After a first step of centrifugation at 400 xg for 8 min at 4°C, dissociated cells were passed through a 70 μm filter, rinsed with complete RPMI and centrifuged at 400 xg for 5 min at 4°C. After a treatment with 5 mL red blood cell lysis (Roche) at 4°C for 5 minutes, cells were washed twice with PBS 2 mM EDTA 0.1% BSA (named FACS buffer) and viable immune cells were enriched in a 40:80% Percoll® gradient.

#### Human tissue dissociation

The tissues were collected in CO_2_ independent medium (Fisher sci., Cat# 11580536) and kept on ice until processing within one hour. One piece was placed in RNA later and conserved at −80°C until RNA extraction. One histological slice was fixed in 10% buffered formalin phosphate (Sigma) and paraffin-embedded. HES-stained slides were reviewed by a pathologist (Figure S1). The other slice was embedded in OCT (VWR) and put in a bath of isopentane placed in a liquid nitrogen bath. The OCT-embedded tissue blocks were stored in a sealed container at −80°C until cryo-sectioning. The last tissue was rinsed in RPMI, transferred to a sterile 60 mm^2^ tissue culture dish and cut in 1 mm^3^ pieces. Tissue pieces were placed in Gentle MACS C tubes and incubated in tissue dissociation solution (tumor dissociation kit, Miltenyi) using the GentleMACS dissociator (program h_TDK_2, Miltenyi). After a first step of centrifugation at 400 xg for 8 min at 4°C, dissociated cells were passed through a 40 μm filter, rinsed with complete RPMI and centrifuged at 400 xg for 5 min at 4°C. After a treatment with 5 mL red blood cell lysis (Roche) at 4°C for 5 minutes, cells were washed twice with PBS 2 mM EDTA 0.1% BSA (named FACS buffer) prior to counting in Trypan blue exclusion dye.

#### Flow cytometry staining

Dissociated cells were incubated with cell surface antibodies in FACS buffer for 25 minutes at 4°C in the dark or for 15 munites at room temperature and washed twice before analysis. For intracellular staining of cytokines, cells were fixed and permeabilized with Fixation/Permeabilization Solution Kit according to manufacturer instructions (BD Biosciences). Viable cells were analyzed based on side scatter and viability dye. Analysis was performed on the BD LSRFortessa. Cells from murine samples were blocked in anti-mouse CD16/32 antibody (1:200e) for 10 min at 4°C prior to addition of cell surface antibodies.

#### Single-cell FACS sorting

For scRNAseq experiments, freshly dissociated single-cell suspensions were sorted based on this gating: FITC-CD45 ^+^, APC.Cy7-panTCRαβ^-^ and PE.Cy7-CD19^-^ cells. For the cytotoxicity experiments, freshly dissociated single-cell suspensions from non-tumoral liver were sorted based on this gating: Viogreen-CD45^+^, AF700- CD19^-^, PE-panTCRαβ ^-^, PE-panTCRγδ ^-^ and APC-CD94^+/-^ FITC-HLA-DR^+/-^ or Viobright 515- CD56^+^. For the immunosuppression experiments, dissociated tumor single-cell suspensions were thawed in complete RPMI medium and incubated for 10 minutes at 37 °C to wash out residual DMSO in media. Cells were pelleted and suspended in cold FACS buffer prior to staining with cell surface antibodies: Viogreen-CD45^+^ APC Vio 770-Lin^-^ (CD3, CD19, CD56) FITC-CD36^+^ PE.Cy7-CD163^+/-^ APC-LOX-1^+/-^.Viable cells were analyzed based on side scatter gates and Sytox blue viability dye 1:5000 (Miltenyi). Cells were sorted using FACS Aria II at 4°C in pre-coated (2 hours at 37°C with PBS 10% FCS) 1.5 mL low binding tubes (Eppendorf) containing either PBS-0.04% pure BSA (for scRNAseq experiments) or complete RPMI (for co-culture experiments). Sorted cells were centrifuged 350 xg at 4°C for 5 minutes and counted with trypan blue dye before being processed.

#### Single cell RNA sequencing Experiment

15.000 single CD45+ panTCRαβ- CD19- innate immunity cells were loaded into a chip to form Gel Bead-in-Emulsion in the Chromium Controller. Single-cell libraries were generated using the Single Cell 3’ reagent Kit v3.1 (10X Genomics) as per the manufacturer’s protocol. cDNA was amplified by 12 PCR cycles and 12 cycles were also performed for library preparation (single index PCR). Libraries were pooled and sequenced on a NovaSeq 6000.

#### Visium Spatial gene expression Processing

OCT-embedded tissue blocks were sectioned at 10 μm thickness using Cryostar NX70 (Leica) and tumoral tissues containing RNA with a RIN ≥ 6.8 were included in the Visium experiments. One section per patient was cut at 10 μm thickness and placed on Visium slide capture areas (10x Genomics). Slides were then processed following the 10x Genomics Visium Spatial protocol according to the manufacturer recommendations. The slides containing tissues were methanol-fixed at −20°C and processed for H&E staining and imaging. The slides were scanned using a Nanozoomer 2.0HT (Hamamatsu Photonics France) using objective UPS APO 20X NA 0.75 combined with an additional lens 1.75X. Virtual slides were acquired with a TDI-3CCD camera. Based on tissue optimization experiments performed on HCC#20, HCC tissues were permeabilized for 12 minutes. cDNA was amplified by 14 PCR cycles and 14 cycles were also performed for library preparation (dual index PCR). Libraries were pooled and sequenced on a NovaSeq 6000.

#### Multiplexed Immunohistological Staining and Imaging

3.5μm-thick formalin-fixed paraffin-embedded tissue sections were placed at 37°C overnight and then for 1h at 56°C. The slides were then deparaffinized in fresh xylene and rehydrated in decreasing concentrations of ethanol. The slides were immersed in AR9 buffer, placed in a jar and heated in a microwave for 90 sec at 1000 mw until the buffer boils, followed by an additional microwave treatment for 15 minutes at 160 mw. Slides were allowed to cool down for 20 minutes at room temperature, rinsed with distilled water and then in Tris-Buffered Saline Tween 0.05 (v/v) (TBST) and blocked in antibody diluent / Block (Akoya) for 10 minutes at room temperature. After primary antibody incubation (HLA-DR, 1:400e 1 hour at 37°C) and washing in TBST, secondary-HRP solution was applied onto the slides for 10 minutes at room temperature. After washing in TBST, Opal 520 (Akoya, 1:100e) solution was applied on each slide at room temperature for 10 minutes. Next, slides were rinsed in TBST and placed in AR9 buffer to perform a second cycle of microwave treatment, blocking, primary antibody incubation (GNLY, 1:1000e overnight at 4°C) and Opal 570 deposition (Akoya, 1:100e). At the end of the cycle, slides were counterstained with spectral DAPI (Akoya) and mounted with Prolong Antifade Mountant (Thermofisher sci.). The microscopy was done in the Bordeaux Imaging Center, a service unit of the CNRS-INSERM and Bordeaux University, member of the national infrastructure France BioImaging supported by the French National Research Agency (ANR-10-INBS-04). The confocal images were taken using objectives ACS APO 40X oil NA 1.15 and the lasers used were diodes laser (405 nm, 488 nm and 561 nm).

#### Co-culture experiments

*For the cytotoxicity experiments*, freshly sorted cells were incubated for 12-14h in round bottom 96-well plates with CFSE-labeled K562 cells at a ratio 5:1 in complete RPMI. Briefly, 1 million K562 cells were incubated with 1 µM CellTrace CFSE dye (ThermoFisher sci.) for 20 minutes at 37°C, cells were then incubated 7 minutes at room temperature with complete RPMI and centrifuged prior to seeding with sorted cells. Cells were simulated or not with IL-2 (Peprotech, 20 ng/ mL) and IL-15 (Peprotech, 50 ng/mL). At the end of the co-culture time, Sytox blue was added to the medium (1:5000) and K562-CFSE+ dead cells were quantified using LSRFortessa. *For the immunosuppression experiments with patients-derived MDSCs*, MDSC cells sorted from HCC surgical resections were pre-incubated for 4 hours in round bottom 96-well plates with a TERT-reactive CD4^+^ T lymphocyte clone at a ratio of 5:1 and then stimulated for 12 hours with UCP4 TERT peptides plus BD Golgi Plug each at 1µg/mL final concentration. For the control, T cells were stimulated with Phorbol 12- myristate 13-acetate at 50 ng/mL and ionomycin at 2 µg/mL (Sigma). *For the immunosuppression experiments with HuMoScs,* total T lymphocytes were purified from healthy donor PBMC from buffy coats by the means of magnetic cell sorting using human pan T-cell isolation kits according to manufacturer’s procedure (Militenyi Biotec; 130-096-535). The obtained T-cells were then activated with anti-CD3/CD28-coated beads (Dynabeads, Life Technologies, 11131D) with or without HuMoSCs at different ratios (T-cell/ HuMoSC ratio = 1:1, 1:4). In both immunosuppression tests, cells were pelleted and suspended in FACS buffer prior to staining with cell surface antibodies (CD4-PE C7 and CD3-APC-Vio770) solution containing fixable viability dye. After washing, fixation and permeabilization, intracellular staining with antibody against IFNγ-APC and TNFα1FITC was performed using BD fixation/Permeabilization solutions as per the manufacturer’s recommendations (BD Biosciences). The % inhibition of T cell activity was calculated as: [(% IFNγ^+^ CD4^+^ T cell or TNFα^+^ CD4^+^ T cell in the positive control stimulation i.e. with the TERT-derived peptide or with anti-CD3/CD28 antibodies) − (% IFNγ^+^ CD4^+^ T cell or TNFα^+^ CD4^+^ T cell in the other tested wells) × 100]/(% IFNγ^+^ CD4^+^ T cell or TNFα^+^ CD4^+^ T cell in the positive control stimulation). In the proliferation assay, T cells were stained with CellTrace Violet Dye (ThermoFisher sci. Cat# C34571) according to the manufacturer’s recommendations and then activated with anti-CD3/CD28-coated beads with or without HuMoSCs (ratio 1:1). After 6 days of co-culture, proliferation of CD3^+^ T cells was analyzed by flow cytometry.

#### Orthotopic tumor injection and tumor growth monitoring

10-week old C57BL/6J male mice were treated with buprenorphine s.c. at 0.1mg/kg 30 min prior to anesthesia with isoflurane (2 L/min oxygen). Laparotomy was done to expose the left lateral liver lobe and 20 µL of Hep55.1C cells (0.25 x 10^6^ cells) /Matrigel (7 mg/mL) suspension was gently injected under the liver capsule. A sterile Gel foam was placed on the needle track for 2 min to prevent leakage of cells. Mice were treated with GF9 or control peptides at 25 mg/kg (in 20% propylene glycol, 10% ethanol and 2% Tween 80, i.p. injection) three times a week from day 7 post inoculation of cells till day 21. Tumor growth was monitored using bioluminescence on isoflurane anesthetized mice.

### 5) BIOINFORMATICS METHODS

#### Single-cell RNA-Seq data processing, quality control and cleaning

A data processing summary is presented in Figure S1c and Table S7. Each of the 10X Chromium single-cell Gene Expression data were pre-processed using CellRanger software 6.0.0, including demultiplexing, reads alignment on human reference genome assembly GRCh38 (refdata-gex- GRCh38-2020-A), barcoding and counting of unique UMI. Raw UMI count matrices were imported in R environment to perform deeper quality control steps to exclude low-quality cells. Cells containing less than 300 or more than 4,500 detected features and cells with more than 40,000 counts were discarded. Moreover, cells expressing more than 12% of mitochondrial genes or more than 15% of ribosomal genes were eliminated. A supplementary criteria was added to exclude stressed cells based on a stress response score(Denisenko et al., 2020). The stress response score was calculated using *AddModuleScore* function from Seurat package (version 4.0.1), and the following list of genes: *FOSB, FOS, JUN, JUNB, JUND, ATF3, EGR1, HSPA1A, HSPA1B, HSP90AB1, HSPA8, HSPB1, IER3, IER2, BTG1, BTG2, DUSP1*.

#### Cell doublet detection and removal

Three different doublet predictions were performed using DoubletFinder (version 2.0.3) (McGinnis et al., 2019), scDblFinder (version 1.7.7) (Germain et al., 2021) and scds (version 1.8.0) (Bais and Kostka, 2020). A consensus method was applied, i.e. a cell was considered as a multiplet and discarded if identified in at least two of the three methods. On average, doublets were estimated around 4%, ranging from 1% (sample with the lowest number of cells) to 5.8%. Cleaning and doublets removal account for 10 to 47% of the cells retrieved with CellRanger. Total recovery ranges from 11 to 49% (Suppl. Table 6).

#### Normalization and Data Integration

The 20 pre-processed scRNA-seq data (10 adjacent non-tumoral and 10 tumoral) were normalized using SCTransform (version 0.3.2) with method = «glmGamPoi» before integration. Functions *PrepSCTIntegration* with 3,000 features, *FindIntegrationAnchors* with dims = 30, reduction = «rpca» and reference (4 samples) options, and *IntegrateData* from Seurat were used to integrate all 20 samples.

#### Unsupervised clustering, dimensionality reduction and Data Visualization

Unsupervised clustering, dimensionality reduction and most visualization were performed with Seurat (version 4.0.1). RunPCA function was used with 30 dimensions. The optimized number of dimensions used for RunTSNE and RunUMAP functions was automatically calculated with an in-house script. *FindNeighbors* and *FindClusters* (res=0.5) function were used to predict the 22 clusters described in the main text. *FindAllMarkers* (min.pct = 0.25, logfc.threshold = 0.25) function was used to identify discriminating features between clusters.

#### Cell type/state annotation

Automatic and manual methods were combined to annotate as precisely as possible cell types or states: (1) expression of canonical markers were visualized and lists of discriminating markers were first manually investigated; (2) lists of markers were downloaded from Panglao database, a score method was applied using AddModuleScore and results were visualized on FeaturePlots; and (3) automatic predictions were done using CellTypist (Domínguez Conde et al., 2021) with the whole dataset and cellKB software (Patil and Patil, 2022) using discriminating lists of markers as input.

#### Identification of signature genes

We defined gene signatures as the minimal number of genes allowing to discriminate a cluster from others. Top 20 discriminating genes were investigated and adjusted as necessary to refine the signatures. To validate signatures, scores were calculated using *AddModuleScore* and visualized using *VlnPlot* and *FeaturePlot* functions from Seurat on the different sets. Signature specificity was calculated as the mean expression of a marker gene in a test cluster divided by the mean expression of the same gene in the cluster that expresses it at its second highest level.

#### Trajectory Inference

Trajectory analyses were performed using monocle (version 2.20.0) and slingshot (version 1.6.1) R packages. A reduced number of cells (n = 300) was used to increase the speed of calculation using subset function from Seurat option downsample. Differentially and temporally expressed genes were identified using differentialGeneTest function with options fullModelFormulaStr = ’∼Cluster’ and fullModelFormulaStr = “∼sm.ns(Pseudotime)”, respectively.

#### Estimation of differentiation potency or signaling entropy

The differentiation potency of single-cells was estimated using the SCENT package (Teschendorff and Enver, 2017), with the provided functional gene network «net13Jun12». Normalized count matrices were log(x + 1) transformed. Signaling Entropy Rates (SR) were calculated using Correlation of Connectome and Transcriptome (CCAT) (Teschendorff et al., 2021) with CompCCAT function, implemented in SCENT. High values indicate higher potency or signaling promiscuity/entropy (capacity to differentiate to different lineages). Furthermore, we predicted differentiation states using CytoTRACE (Gulati et al., 2020). We downsampled MNP sets by using 200 cells per cluster and used the online version of CytoTRACE.

#### Differential Expression Analysis, Gene Ontology and Gene Set Enrichment Analysis

Differential gene expression between two clusters or between two conditions were performed using *FindMarkers* function from Seurat. Gene Ontology analyses were done using topGO (version 2.44.0) and org.Hs.eg.db (version 3.13.0) R packages.

#### Pre-processing and visualization of spatial transcriptomic data

Spatial data were pre- processed and aligned using SpaceRanger software v1.3.0 with the reference human genome GRCh38 (refdata-gex-GRCh38-2020-A) to generate raw UMI count spot matrices. Raw UMI counts were normalized using SCTransform. Dimensionality reduction was performed using classical PCA and clustering was performed using Louvain clustering with *FindNeighbors* and *FindClusters* functions, as before. Signatures were spatially visualized using *AddModuleScore* function. Gene expression visualizations were generated using BBrowser BioTuring (Le *et al*. BioRxiv).

#### Transcriptomic source data of external cohorts

To validate the tissue distribution of MNP subsets and their association with clinical parameters, we interrogated several large cohorts of patients with HCC and other solid tumors. First, to validate the enrichment or depletion of cell types, we compared Non Tumoral (NT) and Primary Solid Tumor (T) samples from five different TCGA cohorts: LIHC, KIRC, LUAD, COAD and READ (https://www.cancer.gov/tcga). Normalized count tables were downloaded using TCGAbiolinks R package (Colaprico et al., 2016) with GDCquery function and the following options : data.category = «Gene expression», data.type = «Gene expression quantification», platform = «Illumina HiSeq», file.type = «normalized_results». Clinical data with patient overall survival durations and tumor grades were downloaded with the GDCquery_clinic function. Supplementary clinical information of 196 patients of the LIHC cohort were retrieved from The Cancer Genome Atlas Research Network (Cancer Genome Atlas Research Network. Electronic address and Cancer Genome Atlas Research, 2017). We also tested MNP subsets associations with patient response to anti-PD-1 in ccRCC (Braun *et al*., 2020), and in metastatic melanoma (Hugo *et al*., 2017).

#### Immuno-deconvolution and cancer genomics

We estimated proportions of scRNA-seq cell subsets in the previously cited external cohorts using CIBERSORTx (Steen et al., 2020). We used our MNP set and a second scRNA-seq set containing 58 cell populations including fibroblasts, stromal cells and adaptive immune cells (Qi et al., 2022). To extract the normalized count matrix, we downsampled the number of barcodes per cell type to 200 for our MNP subsets and 100 cells per cell type for the set from Qi et al. (Qi et al., 2022). We built scRNA-seq signature matrices in CIBERSORTx web server. We imputed cell fractions of each immune set within each cohort described in the previous section, with option S-mode ‘Batch correction’. Parsing and visualizations of the proportions were then realized using tidyverse, reshape2, ggplot2, ggsignif, pheatmap, cowplot and corrplot R packages. We performed survival analyses using survminer and survival R packages. Moreover, we verified CIBERSORTx signatures for each cell-type based on signature scoring from file ‘sig_inferred_refsample.bm.K999.txt’. We extracted top-scored 100 genes per cluster. This allowed us to validate our manual signatures to identify highly specific genes.

### 6) STATISTICAL ANALYSES

Statistical analyses were performed with GraphPad Prism version 9 (GraphPad software) or R version 4.1. The statistical tests used are reported in the Figure Legends. LogFC T/NT ratios were calculated as the Log (cell number in the tumor *RCF + 1) – Log (cell number in the non- tumoral tissue *RCF +1) with RCF corresponding to the recovered cell number in either tissue. Cell abundance correlations were calculated using corrplot (version 0.92) and Hmisc (version 4.6.0) R packages. Correlation matrix was done using *cor* function with pearson method and corrplots were calculated using options order=“hclust”.

